# A hypothalamic circuit links hunger to mesolimbic dopamine to drive feeding

**DOI:** 10.64898/2026.09.10.750730

**Authors:** Sam Z. Bacharach, Katherine A. Zappetti, Laryssa O. Coutinho, Amanda Bacherer, Joseph I. Wahba, Heather M. Schneps, Zhong-Wu Liu, Marcelo O. Dietrich, Amber L. Alhadeff

## Abstract

Internal state profoundly influences motivation, yet how the brain prioritizes behavior according to physiological need remains poorly understood. Here, we reveal a neural mechanism through which hunger selectively enhances dopamine responses to food to drive feeding behavior. We demonstrate that hypothalamic AgRP neurons are necessary and sufficient for hunger-induced amplification of food-evoked dopamine signaling, while their activity suppresses dopamine responses to non-food rewards. AgRP neurons do not directly drive dopamine release; rather, they reduce inhibitory inputs onto midbrain dopamine neurons, gating their responses to food. Remarkably, dopamine signaling specifically during encounters with food is required for subsequent hunger-driven consumption. Mechanistically, we show that NPY-mediated inhibition of paraventricular hypothalamic Y1R neurons links AgRP neuron activity to mesolimbic dopamine signaling. Together, these findings reveal how physiological need selectively shapes reward processing to transform hunger signals into motivated behavior.

## Introduction

Internal physiological state shapes how we perceive and respond to the external world^1^. Hunger provides a powerful example of this: food is more rewarding when we are hungry, motivating food seeking and consumption to restore energy balance. The ability to integrate physiological need with environmental rewards is essential for adaptive behavior, yet the mechanisms through which internal state shapes motivation remain incompletely understood. In particular, the precise neural circuitry through which homeostatic hunger circuits interact with brain reward systems to drive feeding behavior remain elusive.

Within the brain, homeostatic (e.g., hypothalamic^2^) and reward (e.g., midbrain dopamine^3^) systems form critical circuits that control food intake. Indeed, manipulation of hypothalamic agouti-related protein (AgRP)-expressing neurons robustly influences feeding behavior and body weight homeostasis^4–6^. Similarly, ventral tegmental area (VTA) dopamine neurons that project to the nucleus accumbens (NAc) play an important role in food motivation and reward seeking^7–10^. Although these systems have traditionally been studied as parallel and independent processes governing food intake, growing evidence suggests that hunger circuits interact with mesolimbic reward and motivational pathways to shape feeding behavior. For example, in humans, fasting amplifies neural and behavioral reward responses to food and food-related cues^11,12^. In rodents, hunger hormones^13,14^ or AgRP neuron activation^15,16^ amplify phasic dopamine responses to food, which encode the motivational value of food and food-associated cues^17–20^.

While these data suggest a coordinated relationship between AgRP and dopamine neurons in response to food, AgRP neurons lack direct anatomical connections with VTA dopamine neurons in adult mice^21,22^, and the neural circuits that mediate this interaction are unknown. Here, using a combination of slice electrophysiology, in vivo neural activity manipulations and monitoring, functional circuit mapping, and behavioral assays, we identify NPY signaling through Y1 receptors in the paraventricular hypothalamus (PVH) as a mechanism through which AgRP neurons selectively enhance mesolimbic dopamine responses to food and drive feeding behavior. Together, these findings uncover a neural circuit through which homeostatic hunger signals engage the mesolimbic dopamine system, revealing how physiological need selectively shapes reward processing to drive motivated behavior.

## Results

### AgRP neuron activity gates VTA dopamine neurons by reducing inhibitory input

We first asked whether AgRP neuron activity directly engages mesolimbic dopamine signaling or instead gates the system in a way that amplifies dopamine responses to behaviorally relevant stimuli. To do so, we chemogenetically activated AgRP neurons with the excitatory receptor, hM3Dq, while recording dopamine dynamics in the NAc using GRAB-DA2h fiber photometry (**Figure 1A and S1A**). For a functional positive control, we stimulated AgRP neurons with the hM3Dq ligand, clozapine-N-oxide (CNO), to show that this manipulation robustly increases food intake in hM3Dq-, but not control eGFP-, expressing mice (**Figure S1B, S1C**). We also verified that dopamine signaling was similar across experimental and control mice under physiological conditions (**Figure S1D**). AgRP neuron activation did not increase NAc dopamine release in the absence of other stimuli (**Figure 1B, C**), suggesting that activity in AgRP neurons alone does not directly trigger dopamine signaling. Because changes in dopamine signaling may be reflected in discrete phasic dopamine events in addition to changes in average dopamine levels over time^23,24^, we next quantified dopamine transient events in the NAc following AgRP neuron activation. AgRP neuron activation modestly increased the number of NAc dopamine transients (**Figure 1D-F**), suggesting that it does not produce a sustained increase in dopamine signaling, but may shift the system into a more excitable state.

**Figure 1.**
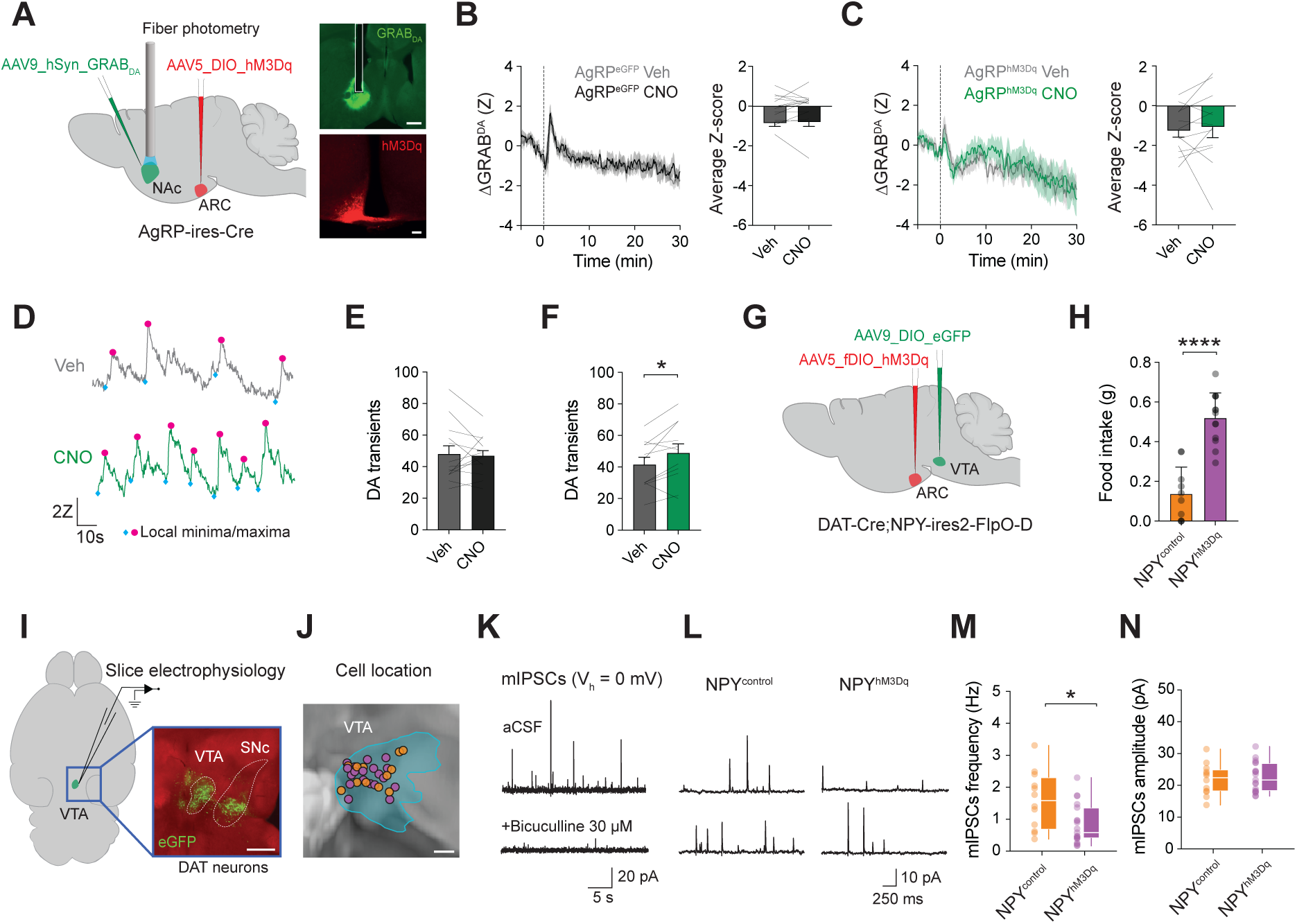
AgRP neuron activity gates VTA dopamine neurons by reducing inhibitory inputs. **(A)** Schematic depicting GRAB-DA fiber photometry recordings in the nucleus accumbens (NAc) during chemogenetic activation of arcuate nucleus (ARC) AgRP neurons. Right, representative images of GRAB-DA in the NAc (scale bar, 500 µm) and hM3Dq in ARC AgRP neurons (scale bar, 100 µm). **(B)** NAc dopamine signals (left) and average Z-score (right) following vehicle (Veh) or clozapine-N-oxide (CNO) administration in control mice (n = 14, paired *t*-test, *p* = 0.7402). **(C)** NAc dopamine signals (left) and average Z-score (right) following Veh or CNO administration in mice expressing hM3Dq in AgRP neurons (n = 12, paired *t*-test, *p* = 0.6418). **(D)** Representative NAc dopamine transients following Veh or CNO administration. **(E)** Number of dopamine transients during the 10 min following Veh or CNO administration in control mice (n = 13, paired *t*-test, *p* = 0.7363). **(F)** Number of dopamine transients during the 10 min following Veh or CNO administration in mice expressing hM3Dq in AgRP neurons (n = 11, paired *t*-test, *p* = 0.0281). **(G)** Schematic depicting chemogenetic activation of NPY/AgRP neurons and labeling of VTA dopamine neurons with eGFP in DAT-Cre;Npy-ires2-FlpO-D mice. **(H)** Food intake in control NPY^control^ mice (n = 8) and NPY^hM3Dq^ mice (n = 13) following chemogenetic activation of NPY/AgRP neurons (unpaired *t*-test, p< 0.0001). **(I)** Schematic depicting ex vivo whole-cell electrophysiological recordings from VTA dopamine neurons following in vivo chemogenetic stimulation of NPY/AgRP neurons. Right, representative image of eGFP labeling in VTA dopamine (DAT-expressing) neurons. Scale bar, 500 µm. **(J)** Anatomical distribution of recorded VTA dopamine neurons (orange cells from control mice, purple cells from experimental mice). Scale bar, 200 µm. **(K)** Representative miniature inhibitory postsynaptic currents (mIPSCs) recorded as outward currents at a holding potential of 0 mV are blocked by the GABAA receptor antagonist bicuculline. **(L)** Representative mIPSCs (two cells per group) from VTA dopamine neurons in NPY^control^ and NPY^hM3Dq^ mice. **(M)** mIPSC frequency in VTA dopamine neurons from NPY^control^ (n = 12 cells from 5 mice) and NPY^hM3Dq^ (n = 19 cells from 7 mice) mice (Mann-Whitney test, U = 54, *p* = 0.01). **(N)** mIPSC amplitude in VTA dopamine neurons from NPY^control^ (n = 12 cells from 5 mice) and NPY^hM3Dq^ (n = 19 cells from 7 mice) mice (Mann-Whitney test, U = 124, *p* = 0.7). Data are presented as mean ± SEM. \**p* < 0.05, \*\**p* < 0.01, \*\*\**p* < 0.001, and \*\*\*\**p* < 0.0001.

Therefore, we directly tested whether AgRP neuron activity alters synaptic input onto VTA dopamine neurons. To do so, we crossed NPY-ires2-FlpO-D mice [to chemogenetically activate AgRP neurons, as they co-express Neuropeptide Y (NPY)^25,26^] with DAT-Cre mice (to label dopamine neurons with eGFP) (**Figure 1G**). After confirming that NPY/AgRP neuron activation increases food intake (**Figure 1H**), we activated NPY/AgRP neurons in vivo and prepared brain sections containing the VTA 60 min later to measure miniature inhibitory and excitatory postsynaptic currents (mIPSCs/mEPSCs) on VTA dopamine neurons (**Figure 1I, J**). For mIPSCs, we verified that outward currents are blocked by the GABAA receptor antagonist bicuculline (**Figure 1K**). NPY/AgRP neuron activation significantly reduced the frequency, without altering the amplitude, of mIPSCs on VTA dopamine neurons (**Figure 1L-N**). In a mixed-effects model accounting for cells nested within mice, the estimated mean frequency was 0.62 Hz in experimental cells and 1.32 Hz in controls, corresponding to a 53% reduction in mIPSC frequency (experimental/control ratio, 0.47; 95% CI, 0.27-0.81; *p* = 0.006; 31 cells from 12 mice) (**Figure 1L-N**). In contrast, AgRP neuron activation did not alter mEPSC frequency or amplitude (**Figure S2**). Together, these results indicate that AgRP neuron activation induces synaptic plasticity onto VTA dopamine neurons by reducing the frequency of inhibitory inputs.

### AgRP neuron activity increases NAc dopamine responses specifically to food

We next probed the involvement of AgRP neurons in stimulating dopamine signaling in response to food and non-food stimuli. We activated AgRP neurons and gave mice access to chow while simultaneously recording NAc dopamine signaling (**Figure 2A, S3A-D**). AgRP neuron activation robustly amplified NAc dopamine signaling evoked by the presentation of food, whereas there were no changes observed in control mice (**Figure 2B, C**)^15^. To test whether this effect depended on prior experience with the food, we presented mice with novel chocolate-flavored grain pellets on two separate days while activating AgRP neurons during both sessions. Dopamine responses were modest during the initial exposure but increased substantially after experience with the food (**Figure 2D, S3E-G**). These findings demonstrate that AgRP neuron activation increases dopamine responses to known foods.

**Figure 2.**
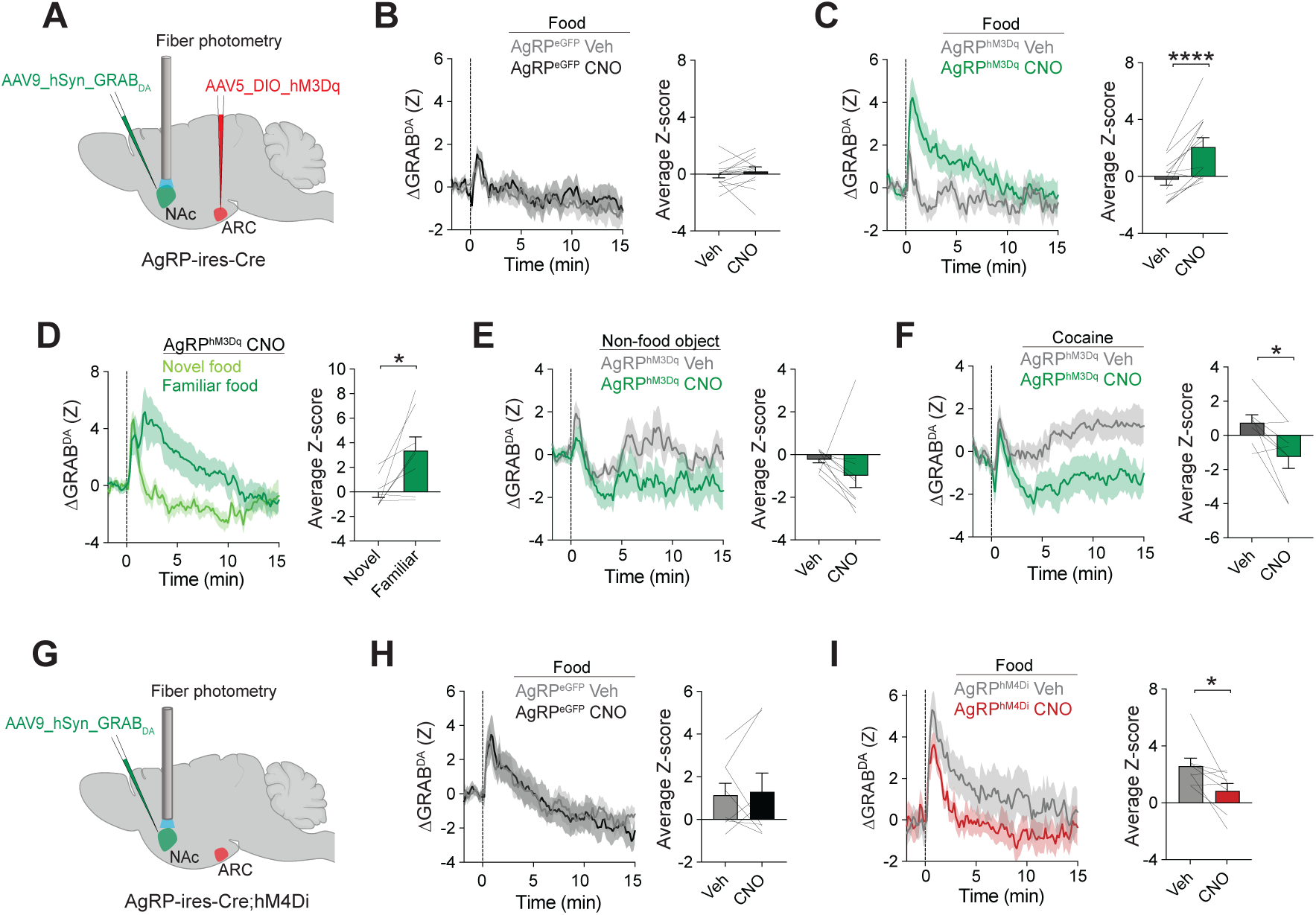
AgRP neuron activity increases NAc dopamine responses specifically to food. **(A)** Schematic depicting GRAB-DA fiber photometry recordings in the nucleus accumbens (NAc) during chemogenetic activation of arcuate nucleus (ARC) AgRP neurons. **(B)** NAc dopamine signals (left) and average Z-score (right) to presentation of food following vehicle (Veh) or Clozapine-N-Oxide (CNO) administration in control mice expressing eGFP in AgRP neurons (n = 14, paired *t*-test, average Z-score, *p* = 0.4397). **(C)** NAc dopamine signals (left) and average Z-score (right) to presentation of food following Veh or CNO administration in mice expressing hM3Dq in AgRP neurons (n = 12, paired *t*-test, average Z-score, *p* < 0.0001). **(D)** NAc dopamine signals (left) and average Z-score (right) to novel and familiar food following AgRP neuron activation (n = 8, paired *t*-test, average Z-score, *p* = 0.0211). **(E)** NAc dopamine signals (left) and average Z-score (right) to presentation of a non-food object following Veh or CNO administration in mice expressing hM3Dq in AgRP neurons (n = 10, paired *t*-test, average Z-score, *p* = 0.2621). **(F)** NAc dopamine signals (left) and average Z-score (right) to 5 mg/kg cocaine following Veh or CNO administration in mice expressing hM3Dq in AgRP neurons (n = 8, paired *t*-test, average Z-score, *p* = 0.0111). **(G)** Schematic depicting GRAB-DA fiber photometry recordings in the NAc during chemogenetic inhibition of ARC AgRP neurons. **(H)** NAc dopamine signals (left) and average Z-score (right) to presentation of food in overnight food-restricted mice following Veh or CNO administration in control mice expressing eGFP in AgRP neurons (n = 8, paired *t*-test, average Z-score, *p* = 0.8147). **(I)** NAc dopamine signals (left) and average Z-score (right) to presentation of food following Veh or CNO administration in overnight food-restricted mice expressing hM4Di in AgRP neurons (n = 8, paired *t*-test, average Z-score, *p* = 0.0321). Data are presented as mean ± SEM. \**p* < 0.05, \*\**p* < 0.01, \*\*\**p* < 0.001, and \*\*\*\**p* < 0.0001.

We next asked whether AgRP neurons broadly amplify dopamine responses to salient stimuli, or instead selectively enhance responses to food. To distinguish these possibilities, we gave mice access to a non-food object (Eppendorf tube) or a highly rewarding drug (cocaine). AgRP neuron stimulation did not potentiate, but actually decreased, dopamine responses to the object (non-significant trend, **Figure 2E, S3H**) and to cocaine (**Figure 2F, S3I**). Thus, AgRP neuron activation does not broadly increase evoked dopamine release. Instead, it differentially modulates dopamine responses according to the nature of the stimulus, enhancing responses to food while suppressing responses to non-food stimuli.

Having demonstrated that AgRP neuron activation selectively enhances dopamine responses to food, we next asked whether endogenous AgRP neuron activity is required for food-evoked dopamine release during hunger. To do so, we chemogenetically inhibited AgRP neurons in food restricted mice and measured NAc dopamine release to food presentation (**Figure 2G**). Inhibition of AgRP neurons markedly suppressed NAc dopamine responses to food (**Figure 2H, I**). Taken together, these results indicate that AgRP neuron activity is both necessary and sufficient for hunger-evoked dopamine responses to food.

### Dopamine release is critical for AgRP neuron-driven feeding behavior

Having established that AgRP neurons selectively amplify dopamine responses to food, we next sought to determine the behavioral significance of this signaling. We hypothesized that the midbrain dopamine system translates the AgRP-mediated hunger signal into a motivational drive for food consumption. If this is the case, then AgRP neuron-evoked dopamine responses to food should relate to subsequent consumption. To examine this, we correlated AgRP neuron-evoked NAc dopamine responses to food with subsequent food intake. Dopamine release following food presentation strongly predicted the amount of food consumed (**Figure 3A**). The early (0-5 min) evoked dopamine response to food was the greatest predictor of future consumption; dopamine signaling in the session had a progressively weaker relationship for predicting consumption (**Figure 3B**). These results suggest that early AgRP neuron-dependent amplification of dopamine responses to food is linked to the magnitude of feeding.

**Figure 3.**
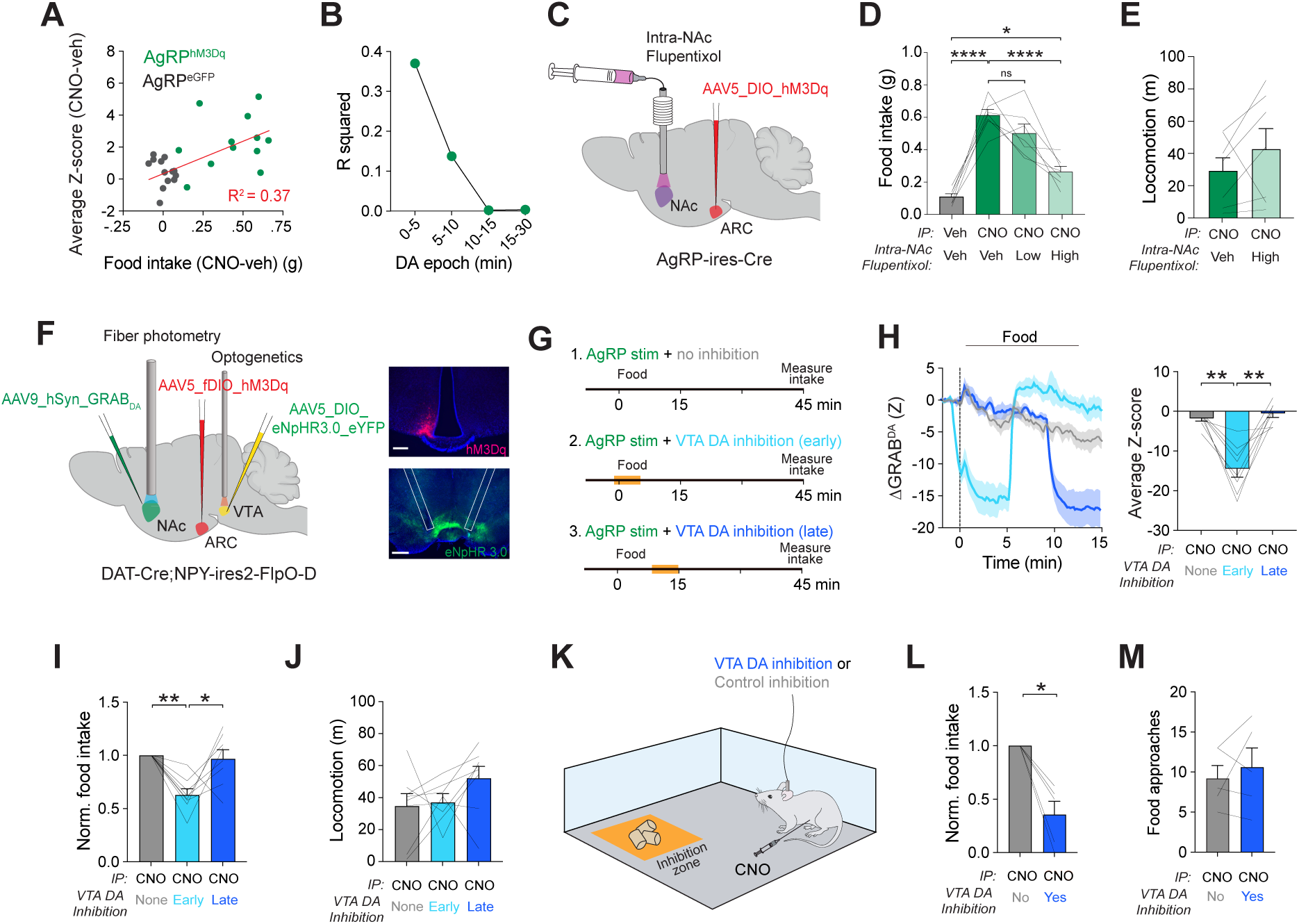
Dopamine release is critical for AgRP neuron-driven feeding behavior. **(A)** Relationship between AgRP neuron stimulation-induced changes in NAc dopamine responses (average Z-score during the first 5 min after food presentation) and subsequent food intake (45 min) in control eGFP mice (gray, n = 14) and mice expressing hM3Dq in AgRP neurons (green, n = 12). Values were calculated as CNO minus vehicle for both measures (Pearson correlation, R² = 0.3705, *p* = 0.0010). **(B)** Relationship between AgRP neuron stimulation-induced changes in NAc dopamine responses during successive post-food epochs and subsequent food intake (45 min) in control eGFP mice (n = 14) and mice expressing hM3Dq in AgRP neurons (n = 12). Values were calculated as CNO minus vehicle (Pearson correlation; 0-5 min, R² = 0.3705, *p* = 0.0010; 5-10 min, R² = 0.1378, *p* = 0.0619; 10-15 min, R² = 0.0023, *p* = 0.8132; 15–30 min, R² = 0.0034, *p* = 0.7758). **(C)** Schematic depicting intra-NAc infusion of the dopamine receptor antagonist flupentixol during chemogenetic activation of ARC AgRP neurons. **(D)** Food intake following i.p. vehicle or CNO administration with intra-NAc vehicle, 12 µg flupentixol, or 20 µg flupentixol (n = 7, repeated-measures ANOVA, *F*(1.578, 9.469) = 42.99, *p* < 0.0001). **(E)** Locomotor activity following AgRP neuron activation with intra-NAc vehicle or 20 µg flupentixol (n = 7, paired *t*-test, *p* = 0.2660). **(F)** Schematic depicting simultaneous GRAB-DA fiber photometry in the NAc, chemogenetic activation of NPY/AgRP neurons, and optogenetic inhibition of VTA dopamine neurons in DAT-Cre;Npy-ires2-FlpO-D mice. Right, representative images of hM3Dq in ARC AgRP neurons (scale bar, 100 µm) and eNpHR3.0 in VTA dopamine neurons (scale bar, 500 µm). **(G)** Experimental timeline for NPY/AgRP neuron activation with (1) no VTA dopamine neuron inhibition, (2) early inhibition following food presentation, or (3) late inhibition following food presentation. **(H)** NAc dopamine signals (left) and average Z-score (first 5 min following food presentation, right) during NPY/AgRP neuron activation with no, early, or late VTA dopamine neuron inhibition (n = 8, repeated-measures ANOVA, *F*(1.298, 9.086) = 26.13, *p* = 0.0004). **(I)** Normalized food intake following NPY/AgRP neuron activation with no, early, or late VTA dopamine neuron inhibition (n = 8, repeated measures ANOVA, *F*(1.910, 13.37) = 9.710, *p* = 0.0027). **(J)** Locomotor activity during the same conditions (n = 8, repeated-measures mixed-effects ANOVA, *F*(1.587, 9.521) = 2.084, *p* = 0.5255). **(K)** Schematic depicting closed-loop optogenetic inhibition of VTA dopamine neurons during food approaches with NPY/AgRP neuron activation. **(L)** Normalized food intake during control or VTA dopamine neuron inhibition trials (n = 5, paired *t*-test, *p* = 0.0331). **(M)** Number of food approaches during control or VTA dopamine neuron inhibition trials (n = 5, paired *t*-test, *p* = 0.5254). Data are presented as mean ± SEM. \**p* < 0.05, \*\**p* < 0.01, \*\*\**p* < 0.001, and \*\*\*\**p* < 0.0001.

We therefore tested whether dopamine signaling within the NAc is required for AgRP neuron-driven feeding. To this end, a dopamine receptor antagonist (flupentixol, broad-spectrum D1 and D2 receptor antagonist) was locally infused into the NAc prior to chemogenetic activation of AgRP neurons (**Figure 3C, S4A**). Blocking dopamine receptors dose-dependently reduced the feeding response normally elicited by AgRP neuron activation (**Figure 3D**).

Importantly, dopamine receptor blockade did not significantly alter locomotor activity (**Figure 3E**), indicating that the reduction in feeding was not due to nonspecific motor impairments.

Because early dopamine signaling was most strongly associated with subsequent intake (**Figure 3B**), we next tested whether dopamine signaling during this early phase is necessary for AgRP neuron-evoked food intake. To do so, we optogenetically inhibited VTA dopamine neurons only during the first five minutes after food presentation after NPY/AgRP neuron stimulation in DAT-Cre;NPY-ires2-FlpO-D mice (**Figure 3F, G, S4B**). This manipulation substantially reduced NAc dopamine signaling only during optogenetic inhibition (**Figure 3H**, quantification from 0-5 min). Strikingly, acute VTA dopamine neuron inhibition resulted in reduced NPY/AgRP neuron-evoked food intake measured over the subsequent 30 minutes (**Figure 3I**). In contrast, in a control condition where VTA dopamine neurons were inhibited from 10-15 min post-food presentation, there was no effect on subsequent food intake (**Figure 3I**). Neither stimulation paradigm significantly changed locomotion (**Figure 3J**).

We next used closed-loop inhibition to determine whether dopamine signaling when mice approach and engage with food is specifically required for NPY/AgRP neuron-evoked feeding (**Figure 3K**). Optogenetic inhibition of VTA dopamine neurons applied only when mice approached food robustly suppressed NPY/AgRP neuron evoked food intake (**Figure 3L**) without altering the total number of food approaches (**Figure 3M**). The preservation of food approaches suggests that mice retain the ability to seek food but were less likely to initiate or sustain consumption when dopamine signaling was disrupted. Thus, AgRP neuron-evoked dopamine signaling during food engagement may be particularly important for converting food-directed motivation into consummatory behavior. Together, these findings indicate that VTA-NAc dopamine signaling during encounters with food is critical for AgRP neuron-driven feeding.

### AgRP neuron projections to the PVH increase dopamine response to food

It is clear that AgRP neurons communicate with dopamine neurons to promote feeding, but there are no direct AgRP projections to either the VTA or the NAc^21,22^. Therefore, we sought to identify the neural circuit(s) through which AgRP neurons influence mesolimbic dopamine signaling. AgRP neurons send dense, direct projections to multiple hypothalamic and extrahypothalamic targets that have been implicated in feeding, arousal, aversion, and motivated behavior^21^. Leveraging this anatomical organization, we used a functional optogenetic projection-mapping strategy to test whether selective stimulation of any of six major AgRP neuron axon projections enhances NAc dopamine release to food. We first verified that, similar to chemogenetic stimulation, optogenetic stimulation [with channelrhodopsin-2 (ChR2)] of all AgRP neurons markedly enhanced food intake and NAc dopamine release to food (**Figure 4A-C, S5A-C**). We next optogenetically stimulated AgRP neuron axon terminals in either the paraventricular hypothalamus (PVH), lateral hypothalamic area (LHA), paraventricular thalamus (PVT), central amygdala (CEA), periaqueductal grey (PAG), or parabrachial nucleus (PBN) (**Figure 4D, S6A-F**), and as a positive control, measured subsequent food intake. As expected^21,27^, AgRP axon terminal stimulation in the LHA, PVH, or PVT enhanced food intake, whereas stimulation of AgRP projections to the CEA, PAG, or PBN did not (**Figure 4E**).

**Figure 4.**
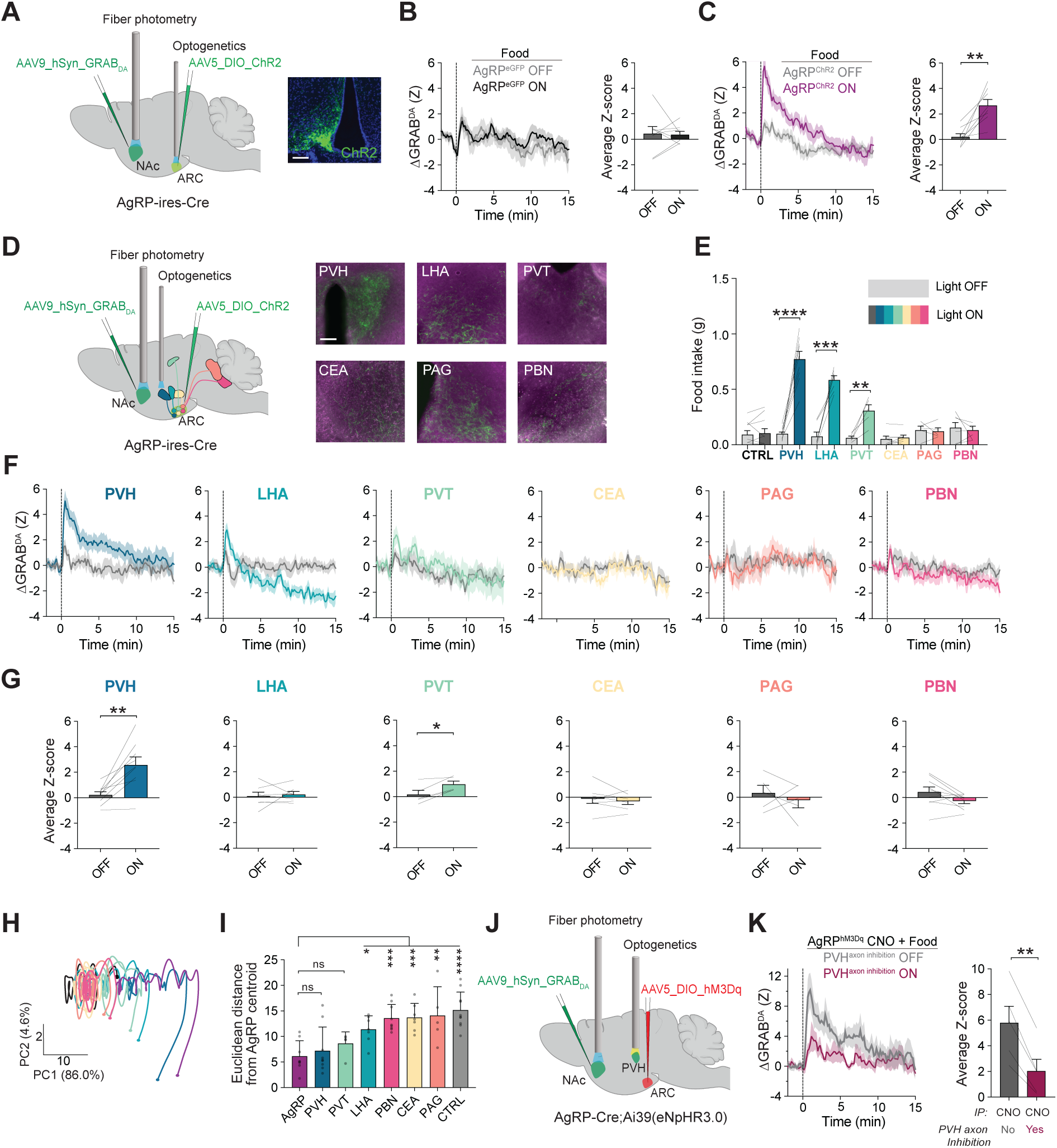
AgRP neuron projections to the PVH increase dopamine responses to food. **(A)** Schematic depicting GRAB-DA fiber photometry recordings in the nucleus accumbens (NAc) during optogenetic stimulation of arcuate nucleus (ARC) AgRP neurons expressing ChR2. Right, representative image of ChR2 in ARC AgRP neurons. Scale bar, 100 µm. **(B)** NAc dopamine signals (left) and average Z-score (right) to food presentation with optogenetic stimulation OFF or ON in control mice expressing eGFP in AgRP neurons (n = 8, paired *t*-test, *p* = 0.9039). **(C)** NAc dopamine signals (left) and average Z-score (right) to food presentation with optogenetic stimulation OFF or ON in mice expressing ChR2 in AgRP neurons (n = 8, paired *t*-test, *p* = 0.0024). **(D)** Schematic depicting NAc GRAB-DA fiber photometry recordings during selective optogenetic stimulation of AgRP neuron projections to either the paraventricular hypothalamus (PVH), lateral hypothalamic area (LHA), paraventricular thalamus (PVT), central amygdala (CEA), periaqueductal gray (PAG), or parabrachial nucleus (PBN). Right, representative images of AgRP axon projections in each target region. Scale bar, 100 µm. **(E)** Food intake with optogenetic stimulation OFF or ON in control mice (n = 10, p = 0.8213) or following stimulation of AgRP projections to the PVH (n = 9, *p* < 0.0001), LHA (n = 7, *p* = 0.0004), PVT (n = 5, *p* = 0.0095), CEA (n = 7, *p* = 0.4021), PAG (n = 5, *p* = 0.7438), or PBN (n = 9, *p* = 0.1842; paired *t*-tests). **(F)** NAc dopamine responses to food presentation with optogenetic stimulation of AgRP projections to the PVH, LHA, PVT, CeA, PAG, or PBN. **(G)** Average NAc dopamine responses (Z) following food presentation with stimulation OFF or ON for AgRP projections to the PVH (n = 9, *p* = 0.0020), LHA (n = 7, *p* = 0.7190), PVT (n = 5, *p* = 0.0137), CEA (n = 7, *p* = 0.4651), PAG (n = 5, *p* = 0.5665), or PBN (n = 9, *p* = 0.0613). All paired *t*-tests. **(H)** Time-resolved principal component analysis (PCA) trajectories of food-evoked NAc dopamine dynamics following AgRP cell-body stimulation or stimulation of individual AgRP neuron projections. Percentages indicate the % total variance explained by each component. **(I)** Euclidean distance of dopamine response dynamics from the AgRP cell-body stimulation centroid across projection stimulation and control conditions (n = 60, one-way ANOVA, *F*(7, 52) = 7.872, *p* < 0.0001). Post hoc comparisons showed that PVH (*p* > 0.9999) and PVT (*p* = > 0.999) stimulation were not significantly different from AgRP cell-body stimulation. **(J)** Schematic depicting NAc GRAB-DA fiber photometry during chemogenetic activation of AgRP neurons and simultaneous optogenetic inhibition of AgRP axon terminals in the PVH. **(K)** NAc dopamine signals (left) and average Z-score (right) to food presentation during AgRP neuron activation with or without inhibition of AgRP axon terminals in the PVH (n = 5, paired *t*-test, *p* = 0.0016). Data are presented as mean ± SEM. \**p* < 0.05, \*\**p* < 0.01, \*\*\**p* < 0.001, and \*\*\*\**p* < 0.0001.

After confirming that baseline dopamine signaling was similar across all groups (**Figure S6G**), we measured the ability of each AgRP neuron projection to potentiate the dopamine response to food. Among all projections, stimulation of AgRP◊PVH axon terminals produced the most robust dopamine response to food (**Figure 4F, G**). AgRP◊PVT axon terminal stimulation modestly increased, whereas projections to the LHA, CEA, PAG, or PBN did not significantly influence, dopamine responses to food (**Figure 4F, G**). Principal component analysis revealed that the dopamine dynamics in response to food produced by AgRP◊PVH stimulation closely matched the effects observed following AgRP neuron cell body activation (more so than all other projections) (**Figure 4H, I**), suggesting the PVH as a key downstream node through which AgRP neurons engage mesolimbic dopamine signaling. Therefore, we tested whether AgRP◊PVH neuron activity is necessary for the feeding and dopaminergic effects of AgRP neuron activation. To do so, we chemogenetically activated all AgRP neurons while simultaneously inhibiting AgRP terminals in the PVH using halorhodopsin (eNpHR3.0) and monitoring food intake **(Figure 4J**). As expected, chemogenetic activation of AgRP neurons increased NAc dopamine release to food and robustly stimulated food intake (**Figure 4K, S6H, I**). Strikingly, inhibition of AgRP axon terminals in the PVH attenuated both the food-evoked dopamine response and subsequent food intake (**Figure 4K, S6H, S6I**). Together, these results demonstrate that the AgRP◊PVH pathway is necessary and sufficient to couple hunger-related hypothalamic activity to food-evoked NAc dopamine release.

### PVH Y1R neuron signaling links AgRP neurons to dopamine release

Having identified AgRP◊PVH projections as mediating the potentiated dopamine response to food by AgRP neurons, we next investigated the local PVH mechanisms underlying this effect. AgRP neurons release multiple inhibitory signals, including NPY, AgRP, and GABA. We therefore asked which AgRP neuron-derived signal(s) acts within the PVH to regulate dopamine release. NPY, SHU9119 (the melanocortin 4 receptor ligand that mimics the antagonistic effect of AgRP), or baclofen (GABAB-receptor agonist) were locally infused into the PVH while NAc dopamine responses to food were measured (**Figure 5A, S7A**). Only NPY infusion significantly enhanced food intake and NAc dopamine release (**Figure 5B, C**). These results suggest that NPY is the principal AgRP neuron-derived signal within the PVH that links hunger to mesolimbic dopamine release.

**Figure 5.**
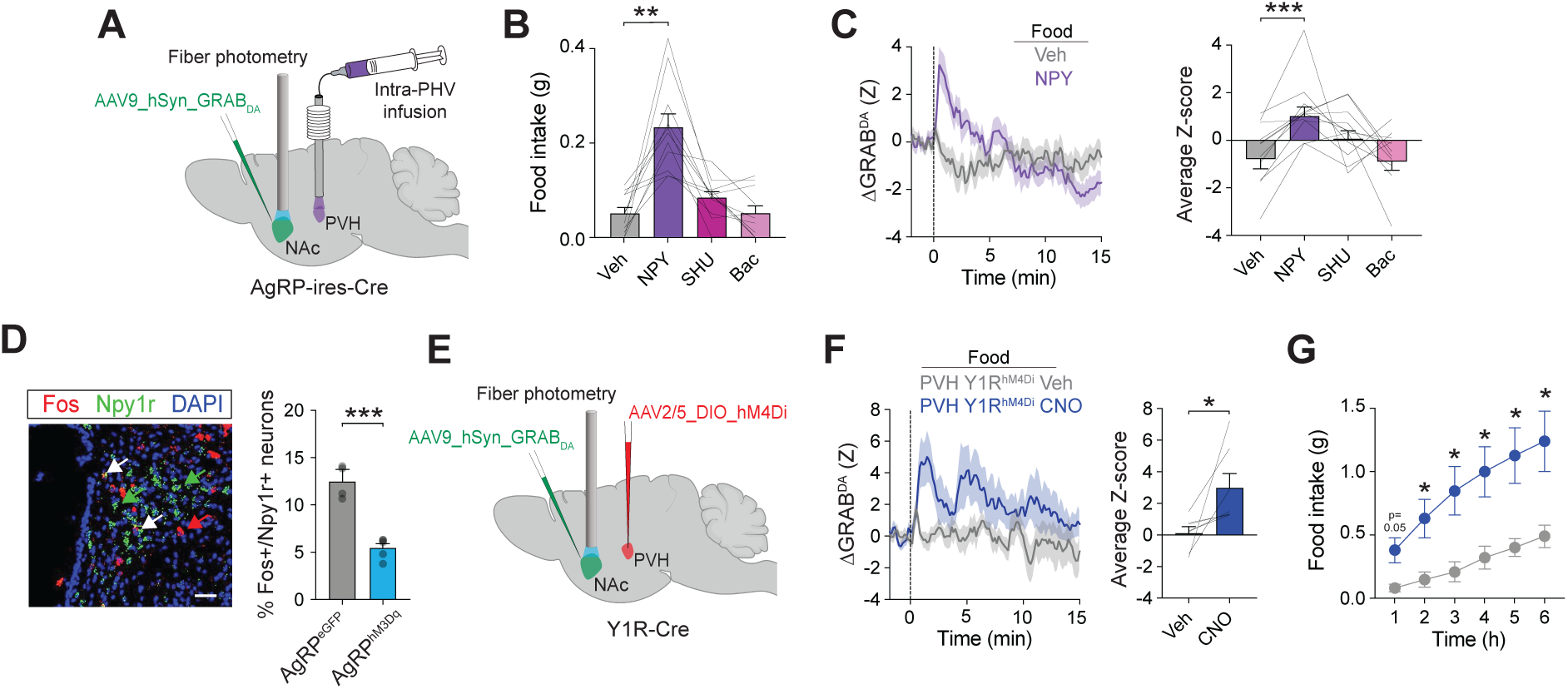
PVH Y1R neuron signaling links AgRP neurons to dopamine release. **(A)** Schematic depicting intra-PVH infusion of AgRP-released neuropeptide and neurotransmitter agonists with simultaneous recording of food-evoked NAc dopamine responses using GRAB-DA fiber photometry. **(B)** Food intake following intra-PVH infusion of vehicle (Veh), NPY, SHU9119 (SHU), or baclofen (Bac) (n = 11, one-way repeated-measures ANOVA, *F*(1.834, 18.34) = 23.91, *p* < 0.0001; Veh vs. NPY, *p* = 0.0013; Veh vs. SHU, *p* = 0.3409 Veh vs. Bac, *p* > 0.9999). **(C)** NAc dopamine signals (left) and average Z-score (right) to food presentation following intra-PVH infusion of Veh, NPY, SHU, or Bac [n = 11, one-way repeated-measures ANOVA, *F*(2.485, 24.85) = 6.108, *p* = 0.0045; Veh vs. NPY, *p* = 0.0007; Veh vs. SHU, *p* = 0.3632; Veh vs. Bac, *p* = 0.9955]. **(D)** Representative image and quantification of Fos expression in Npy1r-expressing PVH neurons following chemogenetic activation of AgRP neurons (eGFP, n = 4; hM3Dq, n = 5; unpaired *t*-test, *p* = 0.0009). Red arrows, Fos+ neurons; green arrows, Npy1r+ neurons; white arrows, co-localized neurons. Scale bar, 50 µm. **(E)** Schematic depicting GRAB-DA fiber photometry recordings in the NAc during chemogenetic inhibition of PVH Y1R neurons in Y1R-Cre mice. **(F)** NAc dopamine signals (left) and average Z-score (right) to food presentation following Veh or CNO administration in mice expressing hM4Di in PVH Y1R neurons (n = 7, paired *t*-test, average Z-score, *p* = 0.0425). **(G)** Cumulative food intake following Veh or CNO administration in mice expressing hM4Di in PVH Y1R neurons (n = 7, two-way repeated-measures ANOVA, main effect of time: *F*(1.237, 7.421) = 35.60, *p* = 0.0003; main effect of treatment: *F*(1.000, 6.000) = 18.96, *p* = 0.0048; time × treatment interaction: *F*(1.612, 9.675) = 9.958, *p* = 0.0059). Data are presented as mean ± SEM. \**p* < 0.05, \*\**p* < 0.01, \*\*\**p* < 0.001, and \*\*\*\**p* < 0.0001.

Because NPY acts through inhibitory Y1 receptors expressed by a population of PVH neurons, we hypothesized that AgRP neurons recruit dopamine signaling by suppressing PVH Y1R neurons. We examined neural activation (i.e. Fos expression) in Y1R-expressing PVH neurons following chemogenetic activation of AgRP neurons. AgRP neuron activation reduced the proportion of Fos+ Y1R-expressing PVH neurons (**Figure 5D**), consistent with AgRP neuron-mediated inhibition of this population.

Finally, we tested whether inhibition of PVH Y1R neurons reproduces the effects of AgRP neuron activation. An inhibitory DREADD, hM4Di, was selectively expressed in PVH Y1R neurons, and NAc dopamine release and food intake were measured following chemogenetic inhibition of this population (**Figure 5E, S7B**). Inhibiting PVH Y1R neurons significantly amplified NAc dopamine release following food presentation and increased food intake (**Figure 5F, G**).

Thus, inhibition of PVH Y1R neurons is sufficient to engage the dopamine and feeding responses elicited by AgRP neuron activation. Together, these findings identify NPY-mediated inhibition of PVH Y1R neurons as a key mechanism through which AgRP neurons recruit mesolimbic dopamine signaling to promote feeding behavior.

## Discussion

Hunger makes food more rewarding, but the underlying neural mechanisms for this remain unclear. Here, we demonstrate that AgRP neurons bidirectionally regulate hunger-induced dopamine release to food, and that this dopamine signaling is causally related to subsequent food intake. Importantly, the potentiated dopamine response during AgRP neuron stimulation is specific to food, as it does not change responses to a non-food object or drug reward. AgRP neurons achieve this effect on dopamine signaling by reducing inhibitory tone onto VTA dopamine neurons, potentially making them more excitable. Mechanistically, the amplified dopamine response to food is mediated by AgRP neuron projections to the PVH, which inhibit Y1R-expressing neurons via NPY signaling. Overall, these findings identify a neural circuit linking homeostatic hunger circuits to increased dopamine signaling, revealing how hunger increases food reward.

We find that AgRP neuron activity potentiates the dopamine response to food, but does not increase tonic NAc dopamine release in the absence of other stimuli. This is consistent with our previous work^15^ and suggests that hunger (via AgRP neuron activation) does not directly activate dopamine neurons, but rather, gates dopamine neurons so they are in a more permissive state. Our electrophysiological data corroborate this: activation of AgRP neurons reduced inhibitory tone onto VTA dopamine neurons. This is also consistent with previous work demonstrating that the hunger hormone ghrelin increases the excitability of VTA dopamine neurons, in part by reducing mIPSC and increasing mEPSC frequency^28^. These findings reconcile two previous results that can seem contradictory: AgRP neuron activity transmits a negative valence signal^29^, but is also positively reinforcing in the context of eating^30^. Our data showing that AgRP neuron activity enhances dopamine signaling only in the context of food explain how these two seemingly opposite findings can simultaneously be true: hunger, as a physiological need state, is intrinsically unpleasant, but at the same time it makes eating much more rewarding, an effect we now show requires AgRP neuron activity.

Although our results resolve the aforementioned paradox, they raise an additional important question: if AgRP neuron activity puts VTA dopamine neurons into a more excitable state, why is the potentiated NAc dopamine signaling specific to food? One possibility is that the dopamine neurons that AgRP neurons influence are “food-specific” neurons. This could be true, as there is some evidence that different subsets of dopamine neurons can respond to different rewards (e.g., water versus nutrients)^31^. Alternatively, AgRP neurons could influence dopamine neurons that are broadly responsive to rewards^32,33^. In this case, hunger/AgRP neuron activity may change how broadly responsive dopamine neurons process incoming sensory information, allowing food-related inputs to produce larger dopamine responses. This possibility fits with evidence that the relatively small population of VTA dopamine neurons support reinforcement across diverse classes of rewarding stimuli, including food, water, specific nutrients, social rewards, alcohol, and drugs (see ^34^ for review).

To determine the circuit mechanisms that mediate the effects of AgRP neurons on dopamine signaling, we systematically screened 6 major AgRP neuron projections. Given that the LHA has a large and well-studied projection to mesolimbic dopamine neurons with established connections to food reward^35–40^, we were surprised that activation of this population had little effect on the dopamine response to food. Instead, our screen revealed AgRP neuron projections to the PVH as the major output influencing food-evoked dopamine signaling. The PVH, and especially the subset of cells that expresses the melanocortin-4 receptor (MC4R), has been classically viewed as a hub for the homeostatic regulation of food intake^41–45^. Our findings add to this literature by showing that the PVH also participates in regulating the motivational value of food. Available transcriptomic and anatomical datasets suggest that Y1R neurons only partially overlap with the MC4R population^46,47^ and other satiety populations such as GLP1R-expressing neurons^47^. Therefore, Y1R neurons (or a subset of them) may represent a unique population of PVH neurons that is involved in reward signaling. Interestingly, a study that combined calcium imaging and RNA multiplexed transcriptional profiling of PVH cell types across 11 behavioral states demonstrated that Y1R neurons were enriched in the hedonic eating state^48^, providing additional support for our finding that the PVH connects homeostatic to reward signaling. It will be important for future studies to determine whether there is a discrete subpopulation of PVH Y1R neurons that mediates the effects of AgRP neuron activity on dopamine responses to food.

Another important question involves exactly how PVH Y1R neurons are communicating with mesolimbic dopamine systems. PVH Y1R neurons directly project to the VTA^47^, providing the most parsimonious connection. However, it is unknown whether they synapse directly with VTA dopamine neurons, GABAergic interneurons, or another cell type. Given that PVH Y1R neurons are primarily glutamatergic^48^, and that AgRP neuron activity reduces inhibitory input onto VTA dopamine neurons (**Figure 1M**), we propose that AgRP neurons inhibit PVH Y1R neurons, reducing excitation of VTA GABAergic neurons that in turn reduce inhibitory tone onto local dopamine neurons. However, it will be necessary to directly test this proposed circuit, as other circuit organizations are certainly possible, such as PVH Y1R neurons projecting to an intermediary inhibitory node (e.g., LDT^49,50^ or peri-LC^51^) that then projects to the VTA.

Our findings link homeostatic and reward systems in the brain, defining key circuitry for motivating feeding behavior when it is most important, i.e., during physiological hunger. In many modern food environments, however, energy-dense foods are plentiful, and therefore extreme hunger may be rare. Previous work suggests that high-fat and high-sugar diets disrupt both AgRP and dopamine neuron signaling^16,52^. Our work connecting these neural populations provides the foundation to understand whether and how this circuit connectivity becomes dysregulated with exposure to highly-palatable foods and/or diet-induced obesity.

Taken together, our data identify the neural circuitry through which hunger selectively recruits mesolimbic dopamine signaling to promote feeding behavior. Rather than globally increasing reward processing, hunger gates dopamine responses specifically to food through an AgRP◊PVH Y1R neuron pathway. Overall, this work establishes a mechanistic understanding of how homeostatic and reward systems interact to drive feeding behavior.

## Resource availability

### Lead contact

Requests for resources or reagents should be directed to and will be fulfilled by the lead contact, Amber L. Alhadeff

### Materials availability

This study did not generate new materials or reagents.

### Data and code availability

All code used in this study will be made publicly available before or at time of publication. Raw data will be made available by request from the lead contact.

## Acknowledgements

We thank E. Moore, G. Montgomery, O. Iyilikci, and H. Haddock-Martinez for technical assistance and A. McKnight for comments on the manuscript. A.L.A. is a New York Stem Cell Foundation – Robertson Investigator and a Pew Biomedical Scholar. This work was supported by the National Institutes of Health (R01DK131558 and DP2AT011965 to A.L.A., T32DC000014 and F32DK137446 to S.Z.B., and instrument award S10OD030354 for the acquisition of the Leica STELLARIS 5 confocal microscope to A.L.A.), the American Heart Association (857082 to A.L.A.), the New York Stem Cell Foundation (to A.L.A.), the Klingenstein Fund and Simons Foundation (to A.L.A.), the Pew Charitable Trusts (to A.L.A.), and the Monell Chemical Senses Center (to A.L.A.).

## Author contributions

S.Z.B., M.O.D., and A.L.A. conceived and designed the experiments; S.Z.B., K.A.Z., L.O.C, A.B., E.M., J.I.W., H.M.S., Z-W.L., M.O.D., and A.L.A. performed experiments, analyzed data, and/or interpreted data; S.Z.B. and A.L.A. wrote the manuscript with comments from all authors.

## Declaration of interests

A.L.A. is a scientific advisory board member for Zealand Pharma. This work is unrelated to the work presented in the current manuscript.

---

**Figure S1.**
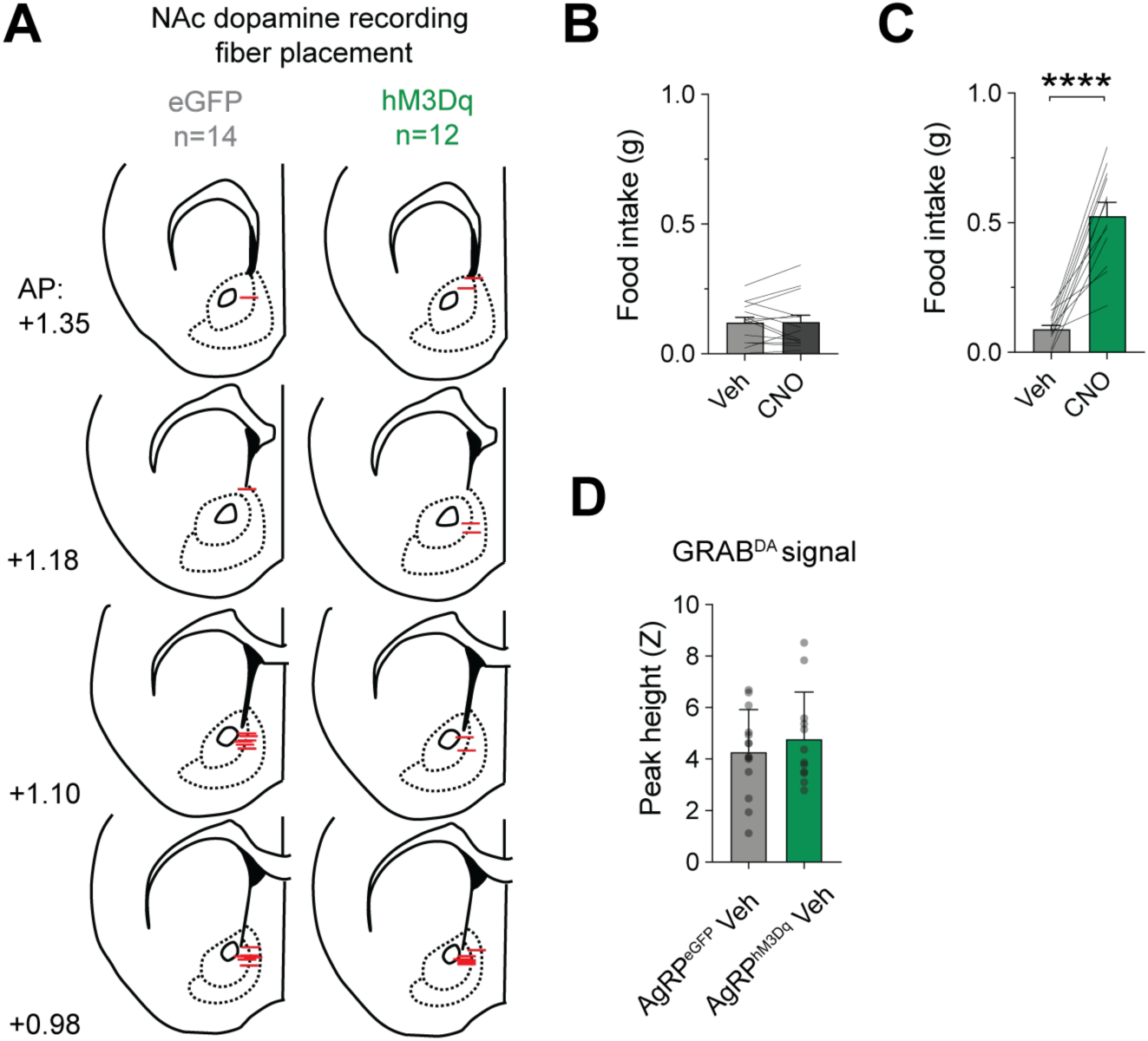
Fiber placements, functional controls, and comparable dopamine responses between AgRP^eGFP^ and AgRP^hM3Dq^ mice. **(A)** Fiber placements in the NAc in control (AgRP^eGFP^) mice (n = 14) and experimental (AgRP^hM3Dq^) mice (n = 12). **(B)** Food intake following vehicle (Veh) or Clozapine-N-Oxide (CNO) administration in AgRP^eGFP^ mice (n = 14, paired *t*-test, *p* = 1.0). **(C)** Food intake following Veh or CNO administration in AgRP^hM3Dq^ mice (n = 12, paired *t*-test, *p* < 0.0001). **(D)** Peak NAc dopamine response in the two minutes following vehicle i.p injection in AgRP^eGFP^ and AgRP^hM3Dq^ mice (eGFP, n = 14; hM3Dq, n = 12; unpaired *t*-test, *p* = 0.4612). Data are presented as mean ± SEM. \**p* < 0.05, \*\**p* < 0.01, \*\*\**p* < 0.001, and \*\*\*\**p* < 0.0001.

**Figure S2.**
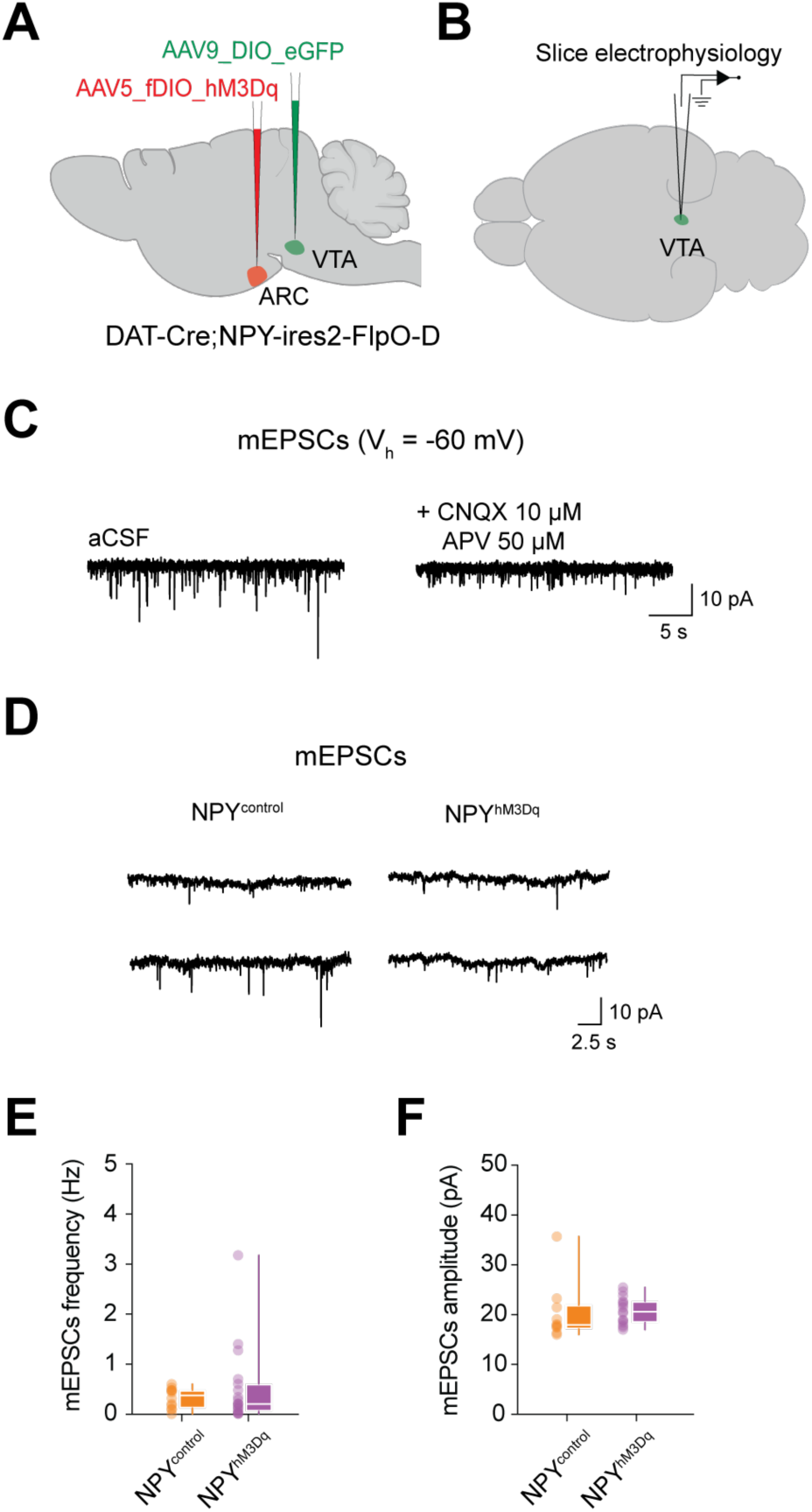
AgRP neuron stimulation does not change mEPSCs on VTA dopamine neurons. **(A)** Schematic depicting chemogenetic activation of NPY/AgRP neurons and labeling of VTA dopamine neurons with eGFP in DAT-Cre;Npy-ires2-FlpO-D mice. **(B)** Schematic depicting ex vivo whole-cell electrophysiological recordings from VTA dopamine neurons following in vivo chemogenetic stimulation of NPY/AgRP neurons. **(C)** Representative miniature excitatory postsynaptic currents (mEPSCs) recorded as inward currents are blocked by the glutamate receptor antagonists CNQX and APV. **(D)** Representative mEPSC (two cells per group) from VTA dopamine neurons in NPY^control^ and NPY^hM3Dq^ mice. **(E)** mEPSC frequency in VTA dopamine neurons from NPY^control^ (n = 12 cells from 6 mice) and NPY^hM3Dq^ (n = 19 cells from 7 mice) mice (Mann-Whitney test, U = 108, *p* = 0.823). **(F)** mEPSC amplitude in VTA dopamine neurons from NPY^control^ (n = 12 cells from 6 mice) and NPY^hM3Dq^ (n = 19 cells from 7 mice) mice (Mann-Whitney test, U = 99, *p* = 0.192). Data are presented as mean ± SEM. \**p* < 0.05, \*\**p* < 0.01, \*\*\**p* < 0.001, and \*\*\*\**p* < 0.0001.

**Figure S3.**
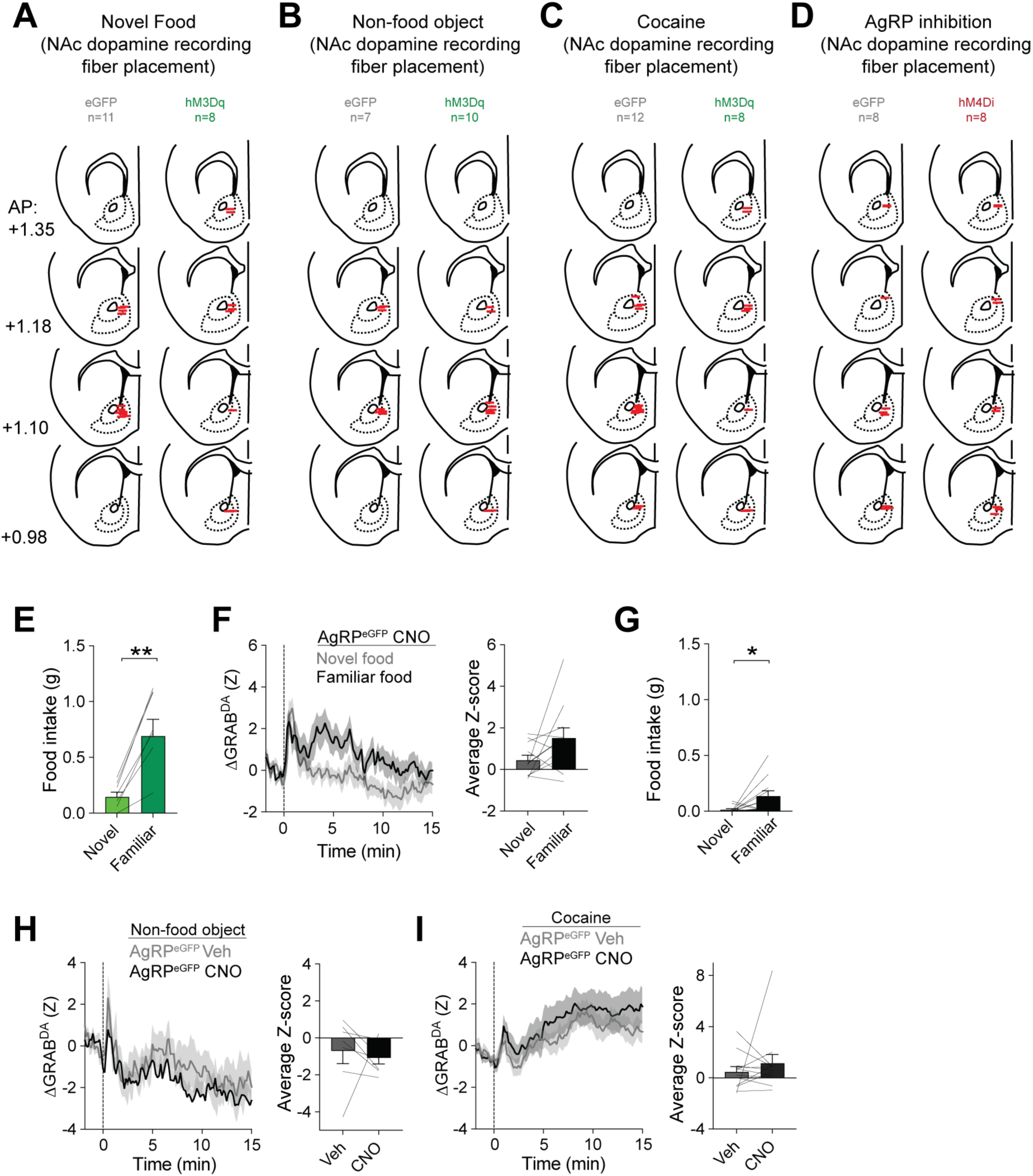
Fiber placements and control measurements for stimulus-evoked NAc dopamine responses. **(A-D)** Fiber photometry placements in the NAc for mice used for novel food **(A)**, non-food object **(B)**, cocaine **(C)**, and AgRP neuron inhibition **(D)** experiments. **(E)** Food intake during exposure to novel food vs. familiar food in AgRP^hM3Dq^ mice (n = 8, paired *t*-test, *p* = 0.0017). **(F)** NAc dopamine signals (left) and average Z-score (right) during the first and second exposure to novel food in control AgRP^eGFP^ mice (n = 11, paired *t*-test, average Z-score, *p* = 0.0831). **(G)** Food intake during exposure to novel food vs. familiar food in AgRP^eGFP^ mice (n = 11, paired *t*-test, *p* = 0.0463). **(H)** NAc dopamine signals (left) and average Z-score (right) to presentation of a non-food object following Veh or CNO administration in control AgRP^eGFP^ mice (n = 7, paired *t*-test, average Z-score, *p* = 0.6814). **(I)** NAc dopamine signals (left) and average Z-score (right) to cocaine following Veh or CNO administration in control AgRP^eGFP^ mice (n = 12, paired *t*-test, average Z-score, *p* = 0.3892). Data are presented as mean ± SEM. \**p* < 0.05, \*\**p* < 0.01, \*\*\**p* < 0.001, and \*\*\*\**p* < 0.0001.

**Figure S4.**
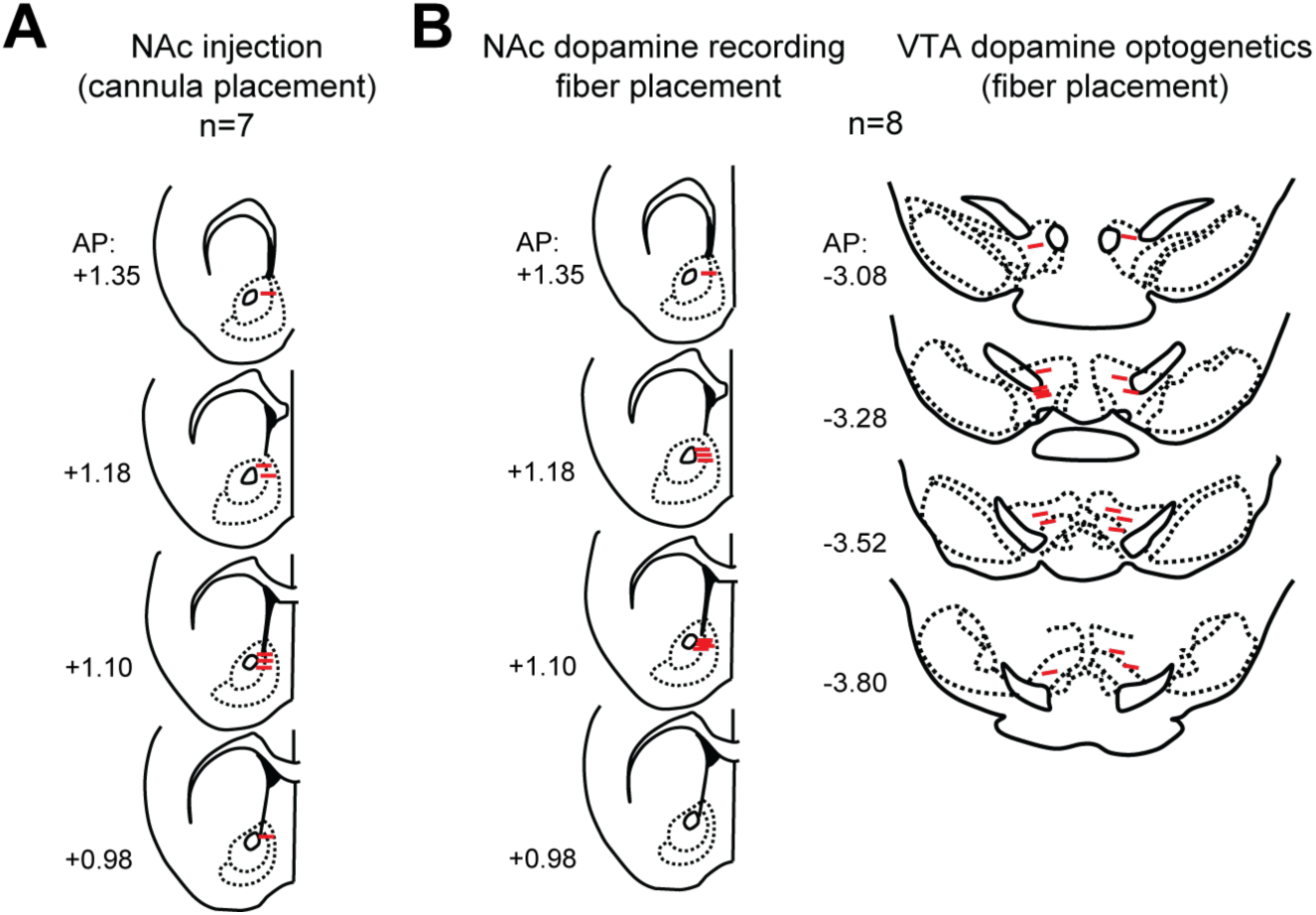
Cannula placements for dopamine antagonist experiments and fiber placements for dopamine inhibition experiments. **(A)** Cannula placements in the NAc for dopamine antagonist experiments. **(B)** Fiber placements in the NAc and optogenetic fiber placements in the VTA for VTA dopamine inhibition experiments.

**Figure S5.**
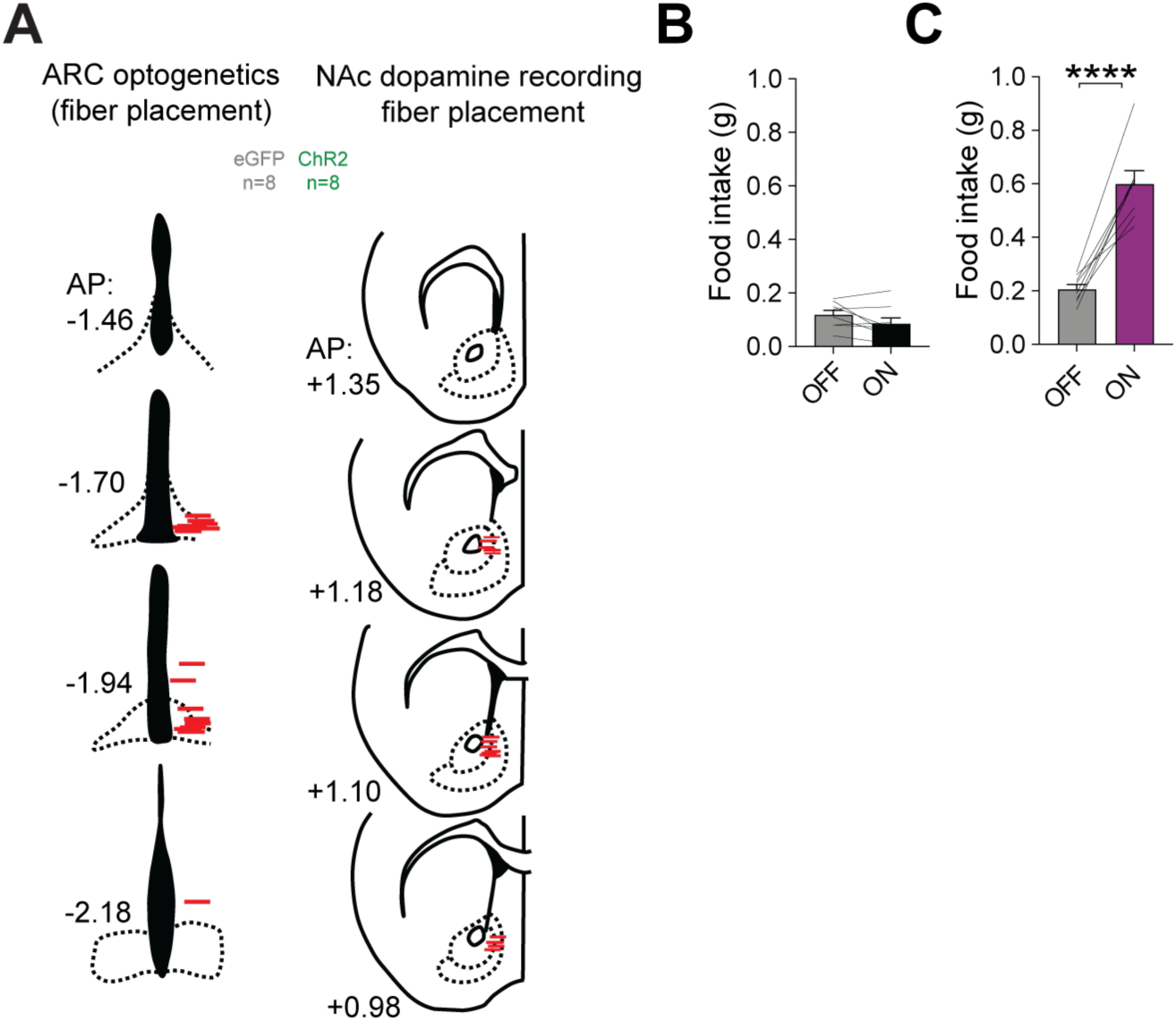
Fiber placements and positive control measurements for optogenetic AgRP neuron stimulation and dopamine recordings. **(A)** Fiber placements for optogenetic stimulation of AgRP neuron cell bodies in the arcuate nucleus (ARC) and GRAB-DA fiber photometry recordings in the nucleus accumbens (NAc). **(B)** Food intake with optogenetic stimulation OFF or ON in control AgRP^eGFP^ mice (n = 8, paired *t*-test, *p* = 0.1263). **(C)** Food intake with optogenetic stimulation OFF or ON in AgRP^ChR^^2^ mice (n = 8, paired *t*-test, *p* < 0.0001). Data are presented as mean ± SEM. \**p* < 0.05, \*\**p* < 0.01, \*\*\**p* < 0.001, and \*\*\*\**p* < 0.0001.

**Figure S6.**
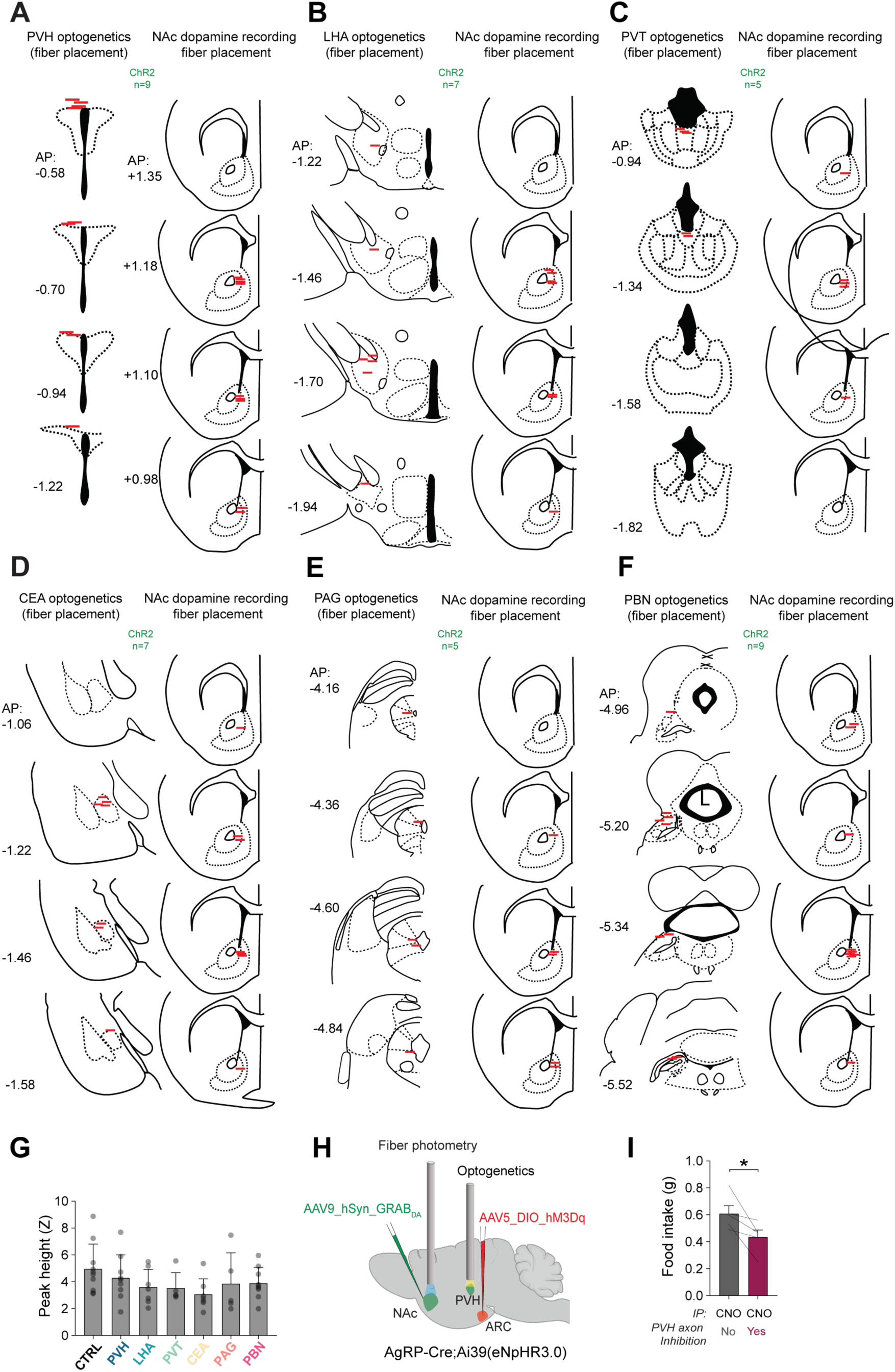
Fiber placements and control data for AgRP axon projection manipulations. **(A-F)** Optogenetic and photometry fiber placements for experiments stimulating AgRP neuron axon projections to the paraventricular hypothalamus (PVH, **A**), lateral hypothalamic area, (LHA, **B**), paraventricular thalamus (PVT, **C**), central amygdala (CEA, **D**), periaqueductal gray (PAG, **E**), and parabrachial nucleus (PBN, **F**). **(G)** Peak NAc dopamine response following vehicle i.p injection in controls and AgRP neuron axon stimulation mice (n = 52, one-way ANOVA, F(6, 45) = 1.125*, p* = 0.3633). **(H)** Schematic depicting NAc GRAB-DA fiber photometry during chemogenetic activation of AgRP neurons and simultaneous optogenetic inhibition of AgRP axon terminals in the PVH. **(I)** Food intake during AgRP neuron activation with or without inhibition of AgRP axon terminals in the PVH (n = 5, paired *t*-test, *p* = 0.0446) Data are presented as mean ± SEM. \**p* < 0.05, \*\**p* < 0.01, \*\*\**p* < 0.001, and \*\*\*\**p* < 0.0001.

**Figure S7.**
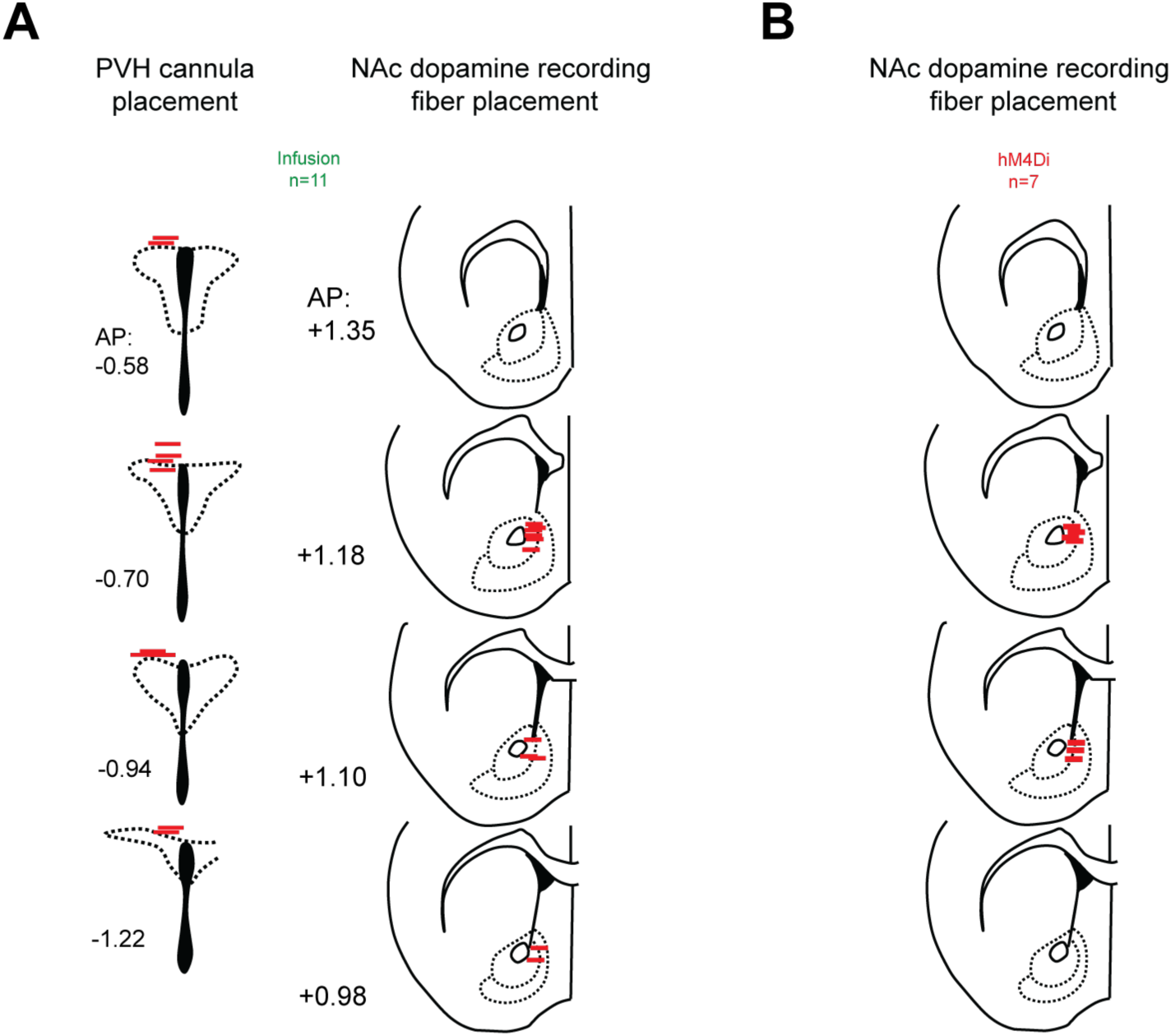
Cannula and dopamine fiber photometry placements for PVH manipulation experiments. **(A)** PVH cannula placements and NAc photometry fiber placements for PVH pharmacology experiments. **(B)** NAc photometry fiber placements for PVH Y1R neuron inhibition experiments.

## STAR Methods

### Key Resources Table

| REAGENT or RESOURCE | SOURCE | IDENTIFIER |
| --- | --- | --- |
| **Bacterial and virus strains** |  |  |
| AAV9-hSyn-GRAB-DA2h | Addgene | #140554-AAV9 |
| AAV5-hSyn-FLEX-hM3Dq-mCherry | Addgene | #44361-AAV5 |
| AAV9-hSyn-DIO-eGFP | Addgene | #50457-AAV9 |
| AAV2/5-hSyn-DIO-hM4D(Gi)-mCherry | Addgene | #44362-AAV2<br>#44362-AAV5 |
| AAV5-EF1a-DIO-hChr2(H134R)-EYFP-WPRE-HGHpA | Addgene | #20298-AAV5 |
| AAV5-hSyn-DIO-eNpHR3.0-EYFP | Addgene | #26966-AAV5 |
| AAV8-hSyn-fDIO-hM3D(Gq)-mCherry-WPREpA | Addgene | #154868-AAV8 |
| **Chemicals, peptides, and recombinant proteins** |  |  |
| Clozapine-N-oxide (CNO) | Tocris Bioscience | Cat#4936 |
| Neuropeptide Y (NPY) | Cayman Chemical | Cat#15071 |
| SHU9119 | Tocris Bioscience | Cat#3420 |
| (RS)-Baclofen | Tocris Bioscience | Cat#0417 |
| Flupentixol dihydrochloride | TargetMol | Cat#T5135 |
| Cocaine hydrochloride | Sigma-Aldrich | Cat#C5776 |
| Tetrodotoxin (TTX) | Sigma-Aldrich | Cat#554412 |
| CNQX | Sigma-Aldrich | Cat#504914 |
| APV (D-AP5) | Cayman Chemical | Cat#14540 |
| Bicuculline | Sigma-Aldrich | Cat#14340 |
| VECTASHIELD antifade mounting medium with DAPI | Vector Laboratories | Cat#H-1200-10 |
| Mowiol 4-88 mounting medium | Sigma-Aldrich | Cat#81381 |
| Chocolate-flavored grain pellets (5TUL) | TestDiet | Cat#181123 |
| Standard rodent chow (LabDiet/Purina 5001) | Purina / LabDiet | Product#5001 |
| Meloxicam | Covetrus | 6451603845 |
| Bupivacaine | Moore Medical | 52683 |
| Isoflurane (anesthetic) | Clipper | 0010250 |
| **Critical commercial assays** |  |  |
| RNAscope Multiplex Fluorescent Reagent Kit v2 | Advanced Cell Diagnostics / Bio-Techne | Cat#323100 |
| **Experimental models: Organisms/strains** |  |  |
| Mouse: C57BL/6J | The Jackson Laboratory | Cat#000664;<br>RRID:IMSR_JAX:000664 |
| Mouse: Agrp-Ires-Cre | The Jackson Laboratory | Cat#012899;<br>RRID:IMSR_JAX:012899 |
| Mouse: Ai39 (RCL-eNpHR3.0/EYFP) | The Jackson Laboratory | Cat#014539;<br>RRID:IMSR_JAX:014539 |
| Mouse: Npy-ires2-FlpO-D | The Jackson Laboratory | Cat#030211;<br>RRID:IMSR_JAX:030211 |
| Mouse: Npy1r-Cre/GFP | The Jackson Laboratory | Cat#030544;<br>RRID:IMSR_JAX:030544 |
| Mouse: Dat-ires-Cre | The Jackson Laboratory | Cat#006660;<br>RRID:IMSR_JAX:006660 |
| Mouse: R26-LSL-Gi-DREADD (hM4Di) | The Jackson Laboratory | Cat#026219;<br>RRID:IMSR_JAX:026219 |
| **Oligonucleotides** |  |  |
| RNAscope probe: Mm-Fos | Advanced Cell Diagnostics | Cat#316921 |
| RNAscope probe: Mm-Npy1r | Advanced Cell Diagnostics | Cat#427021 |
| **Software and algorithms** |  |  |
| MATLAB (R2026a) | MathWorks | https://www.mathworks.com/products/matlab.html;<br>RRID:SCR_001622 |
| GraphPad Prism 11 | GraphPad Software | https://www.graphpad.com/scientific-software/prism/;<br>RRID:SCR_002798 |
| TDT Synapse | Tucker-Davis Technologies | https://www.tdt.com/software/synapse/ |
| AxoGraph X | AxoGraph Scientific | https://axograph.com/ |
| Fiji/ImageJ | NIH / Fiji | https://fiji.sc/;<br>RRID:SCR_002285 |
| **Other** |  |  |
| Fiber photometry optical fiber: 400-um core, NA 0.67 | Doric Lenses | Model MF2.5,<br>400/430-0.67 |
| Optogenetic optical fiber: 200-um core, NA 0.37 | Doric Lenses | Model MFP_200/220/900-0.37 |
| Optogenetic fiber rotary joint | Doric Lenses | Model FRJ_1x1_FC-FC |
| Guide cannula, 26G | RWD Life Science | Model 62003 |
| Internal cannula/injector, 33G | RWD Life Science | Model 62204 |
| 450-nm optogenetic laser system | Lasever | Model LRS450NL-1W-FC+LSR-PS-II |
| RZ10x photometry processor | Tucker-Davis Technologies | Model RZ10x |
| MultiClamp 700B amplifier | Molecular Devices | Model MultiClamp 700B |
| BZ-X800 microscope/slide scanner | Keyence | Model BZ-X800 |
| Stellaris 5 confocal microscope | Leica Microsystems | Model STELLARIS 5 |
| Intracranial cannula fixing screw | RWD Life Science | Model 62502 |
| Metabond cement | Parkell | Cat#S380 |
| Ortho-Jet BCA liquid (dental acrylic) | Lang Dental Manufacturing | Cat#B1306 |
| Jet Tooth Shade Powder (dental acrylic) | Lang Dental Manufacturing | Cat#143069 |

### Method Details

#### Experimental subjects

C57BL/6J (JAX #000664), AgRP-ires-Cre (JAX #012899), Ai39 (eNpHR3.0) (JAX #014539), Npy-ires2-FlpO-D (JAX #030211), Npy1r(Y1R)-Cre/GFP (JAX #030544), Dat-ires-Cre (JAX #006660), and R26-LSL-Gi-DREADD (hM4Di) (JAX #026219) mice were obtained from The Jackson Laboratory and bred for use in experiments. AgRP-Cre x Ai39, AgRP-Cre x hM4Di, Npy-Flp x Npy1r-Cre, and Npy-Flp x Dat-Cre crosses were generated in-house from respective founder lines.

Mice were group-housed under a 12-h light/12-h dark cycle with ad libitum access to standard laboratory chow (Purina 5001) and tap water unless otherwise noted. Adult male and female mice (≥8 weeks of age) were used for experiments, except for slice electrophysiology experiments where mice underwent surgery at approximately P21-P28 and recordings were performed at P45-50. To assess potential sex differences, data from male and female mice were compared across experimental groups. No significant sex differences were detected, therefore data from both sexes were pooled across all analyses. Mice were habituated to handling and experimental procedures before testing. Animals were randomly assigned to experimental conditions, and conditions for within-subject experiments were counterbalanced. All procedures were approved by the Monell Chemical Senses Center or Yale University Institutional Animal Care and Use Committees.

#### Surgical procedures

Before surgery, mice received subcutaneous meloxicam (5 mg/kg) and bupivacaine (2 mg/kg) for analgesia. Mice were anesthetized in an induction chamber with aerosolized isoflurane (2-3%) and transferred to a stereotaxic apparatus. Viral vectors were delivered using a Harvard Apparatus syringe pump connected to a pulled glass micropipette at a rate of 150-300 nL/min. Following viral infusion, pipettes were left in place for 5-10 min, raised 0.1-0.2 mm and left in place for an additional 2 min, and then slowly withdrawn.

Optical fibers and guide cannulae were secured to the skull using jeweler’s screws, Metabond cement (Parkell, S380), and dental acrylic (Lang Dental Manufacturing; Ortho-Jet BCA liquid, B1306; Jet Tooth Shade Powder, 143069). Mice were allowed at least 2 weeks of recovery before behavioral testing. All coordinates were referenced to bregma at skull surface using Franklin and Paxinos Mouse Brain Atlas (2007).

##### Fiber photometry surgery

For fiber photometry recordings of dopamine signaling in the nucleus accumbens (NAc), 250 nL of AAV9-hSyn-GRAB-DA2h (Addgene 140554-AAV9) was injected unilaterally into the NAc (+1.35 mm AP, +0.80 mm ML, −4.40 mm DV). A ferrule-capped optical fiber (400-µm core, NA 0.67; Doric Lenses, MF2.5, 400/430-0.67) was implanted 0.20 mm dorsal to the viral injection site in the same surgery.

##### Cannulation surgery

For intracranial pharmacology experiments, unilateral guide cannulae (26G, 0.48-mm OD; RWD Life Science, model 62003) were implanted targeting either the nucleus accumbens (NAc) or paraventricular hypothalamus (PVH). NAc cannulae were implanted at +1.35 mm AP, +0.80 mm ML, and −3.40 mm DV. PVH cannulae were implanted at 16° anterior to posterior; -2.55 mm AP, 0.25 mm ML, -4.50 mm DV. Guide cannulae were secured to the skull as described above and fitted with dummy cannulae between experimental sessions. For intracranial infusions, a 33G injector (0.21-mm OD; RWD Life Science, model 62204) extending 1 mm beyond the tip of the guide cannula was inserted through the guide.

##### Chemogenetic surgery

For chemogenetic activation of AgRP neurons, a total of 400 nL of AAV5-hSyn-FLEX-hM3Dq-mCherry (Addgene, 44361-AAV5) or AAV5/9_syn_DIO_eGFP (Addgene, #50457-AAV9) was injected into the arcuate nucleus (ARC) of AgRP-ires-Cre mice at −1.35 mm AP, ±0.25 mm ML, −6.00 and -5.85 mm DV. For chemogenetic inhibition of AgRP neurons, AgRP-Cre and hM4Di mice were crossed so that hM4Di was expressed specifically in AgRP neurons in offspring. For chemogenetic inhibition of PVH Y1R neurons, 75 nL of AAV2/5-hsyn-DIO-hm4d(GI)-mCherry (Addgene, 44362-AAV5) was injected into the PVH of Npy1r-Cre mice at −2.20 mm AP, ±0.20 mm ML, −5.10 mm DV at a 16° anterior to posterior angle.

##### Optogenetic excitation of AgRP neurons and projections

For optogenetic excitation of AgRP cell bodies or axon terminals, 400 nL of AAV5-EF1α-DIO-hChR2(H134R)-EYFP-WPRE-HGHpA (Addgene, 20298-AAV5) or AAV9_syn_DIO_eGFP (Addgene, #50457-AAV9) was injected unilaterally into the ARC of AgRP-ires-Cre mice (-1.35 mm AP, +0.25 mm ML, -6.00 and -5.85 mm DV). Optical fibers were implanted targeting the ARC, PVH, LHA, PVT, CEA, PAG, or PBN as appropriate for each experiment. Fiber specifications, approach angles, and stereotaxic coordinates were as follows: Arc: (16°; -3.20 mm AP, +0.25 mm ML, -6.00 mm DV), PVH (16°; -2.20 mm AP, +0.20 mm ML, -5.00 mm DV), PVT (16°; -2.30 mm AP, +0.20 mm ML, -2.80 mm DV), LHA (16°; -3.00 mm AP, +1.00 mm ML, -5.00 mm DV), CEA (16°; -2.60 mm AP, +3.20 mm ML, -4.40 mm DV), PAG (0°; -4.50 mm AP, +0.20 mm ML, -2.40 mm DV), and PBN (0°; -5.50 mm AP, +1.25 mm ML, -3.10 mm DV).

##### Optogenetic inhibition of AgRP terminals in the PVH

AgRP-Cre and Ai39 mice were crossed so that eNpHR3.0 was expressed specifically in AgRP neurons in offspring. These mice received a 400-nL injection of AAV5-hSyn-FLEX-hM3Dq-mCherry (Addgene, 44361-AAV5) into the ARC to permit simultaneous chemogenetic activation of AgRP neurons. An optical fiber was implanted above the PVH to inhibit AgRP axon terminals using the same coordinates as described above.

##### Optogenetic inhibition of VTA dopamine neurons

Npy-ires2-FlpO-D x Dat-Cre mice received bilateral VTA injections of AAV5-hSyn-DIO-eNpHR3.0-EYFP (250 nL per side, Addgene, 26966-AAV5) into the VTA followed by bilateral optical fiber implantation. Viral injection and fibers were inserted at a 10° lateral to medial angle at -3.2 mm AP, 0.6 mm ML, and -4.4 mm DV.

#### Drugs

Clozapine-N-oxide (CNO; Tocris Bioscience, catalog #4936) was prepared in 2% DMSO and 98% sterile saline. CNO was stored in frozen aliquots and thawed and diluted on the day of testing. CNO was administered i.p. at 1 mg/kg. Vehicle consisted of 2% DMSO and 98% sterile saline.

Neuropeptide Y (NPY; Cayman Chemical, catalog #15071) and SHU9119 (Tocris Bioscience, catalog #3420) were dissolved in sterile saline. (RS)-baclofen (Tocris Bioscience, catalog #0417), and flupentixol dihydrochloride (TargetMol, catalog #T5135) were also dissolved in sterile saline. Drug solutions were stored in frozen aliquots and thawed and diluted on the day of use.

For intra-PVH administration, NPY was administered at 1 µg/200 nL^27,53–55^, SHU9119 at 37 pmol/200 nL^27,55,56^, and baclofen at 20 ng/200 nL^27,55,57^. PBS served as the intra-PVH vehicle. For intra-NAc administration, flupentixol was administered at 12 or 20 µg in 350 nL^20,58^; sterile saline served as vehicle.

Cocaine hydrochloride (Sigma-Aldrich, catalog #C5776) was dissolved in sterile saline and administered i.p. at 5 mg/kg.

#### Food deprivation and restriction

Unless otherwise stated, mice were tested under ad libitum-fed conditions. For experiments performed following food deprivation, food was removed from the home cage 20 h before the start of the experiment. For experiments requiring chronic food restriction, mice were singly housed, weighed at approximately the same time each day, and provided 1.5-2.5 g of chow per day to maintain 85-90% of their free-feeding body weight.

#### Optogenetic stimulation

Optogenetic stimulation of AgRP neurons was performed as we have previously published^59,60^. An Arduino generated TTL pulse trains from a 1-W 450-nm laser (Lasever, LRS450NL-1W-FC+LSR-PS-II) coupled via a rotary joint (Doric, FRJ_1x1_FC-FC) to an optical fiber (200-µm core, NA 0.37, Doric, MFP_200/220/900-0.37). For optogenetic excitation, 450-nm light was delivered at an estimated power of 8-10 mW measured at the tip of the implanted optical fiber. Light was delivered as 10-ms pulses at 20 Hz in 2-s stimulation epochs separated by 3-s periods without stimulation. Optogenetic stimulation was initiated at the designated experimental time point and continued for the remainder of the session. For optogenetic inhibition, continuous 590-nm light was delivered at an estimated power of 12-16 mW measured at the fiber tip and remained on for the duration of the designated inhibition period.

#### Dual-wavelength in vivo fiber photometry

Fiber photometry procedures were performed as previously published^27,60–62^. Fiber photometry was performed using a TDT RZ10x real-time processor and Synapse software (Tucker-Davis Technologies). Mice were tested in clean cages identical in size and configuration to their home cages and containing fresh bedding. Animals were habituated to photometry patch cord tethering before testing.

For GRAB-DA2h recordings, the 465-nm signal channel was modulated at 330 Hz and the 405-nm isosbestic/reference channel was modulated at 210 Hz. Combined excitation light was delivered through a 400-µm core, 0.67 NA low-fluorescence optical fiber patch cord. Emitted fluorescence was collected through the same patch cord, detected using a femtowatt photoreceiver, and demodulated by the RZ10x processor. Data were acquired at 1017 Hz and subsequently downsampled to 10.17 Hz for plotting purposes.

#### Fiber photometry data analysis

##### GRAB-DA analysis

Photometry data were analyzed using custom MATLAB scripts. For GRAB-DA experiments, all reported analyses were performed using the 465-nm signal. The 465-nm signal was analyzed without additional filtering or smoothing. For each animal, the 465-nm signal was converted to a z-score relative to a defined pre-event baseline: Z = (F465 – μ baseline)/σ baseline, where μ baseline and σ baseline represent the mean and standard deviation of the baseline fluorescence signal, respectively. A 2-min baseline was used for analyses spanning 5-15 min, whereas a 5-min baseline was used for longer analyses spanning 15-30 min.

Average Z-score was calculated by taking the average of the Z-scored trace over time for each mouse. For chow, novel food, and object presentation experiments, average Z-score was calculated over the 5-min interval immediately following stimulus presentation. For analyses examining responses to CNO, vehicle, or cocaine injection, average Z-score was calculated over the 30-min post-injection period. Bolded lines on traces represent averages across mice; shaded error on traces represents the SEM across mice.

##### Dopamine transient analysis

Dopamine transient analyses were performed on the 465-nm GRAB-DA signal, similar to ^63^. Transients were quantified during a 10-min interval following CNO or vehicle administration. The 465-nm signal was first detrended using a second-order high-pass Butterworth filter with a cutoff frequency of 0.01 Hz. The detrended signal was then converted to a sliding z-score using a local ±30-s window. Candidate transients were identified using the MATLAB ‘findpeaks’ function, with a minimum inter-peak distance of 2 s. For each candidate peak, the local minimum within the preceding 2 s was identified. A transient was considered valid when the peak rose by at least 2 local z-score units above this preceding local minimum.

##### Dopamine peak analysis

To verify that dopamine responses were comparable across experimental groups, peak dopamine responses were quantified from the Z-scored 465-nm GRAB-DA fluorescence signal. The maximum peak amplitude was identified within the 2-min interval following CNO or vehicle injection using the MATLAB ’findpeaks’ function.

##### PCA analysis

To compare the temporal structure of post-food dopamine responses across AgRP cell body stimulation and AgRP projection target stimulation groups, fiber photometry signals were analyzed using a time-delay principal component analysis in MATLAB. For each animal, a 20-s sliding window was advanced across the post-food trace in 1-s increments. Each window therefore represented the recent temporal history of the dopamine signal at a given point in the response. Sliding windows from all animals and time points were combined into a single matrix and subjected to PCA using the same covariance-based eigendecomposition procedure. PC1 and PC2 scores were then reorganized by animal and time to generate a continuous trajectory for each animal through principal-component space. Group trajectories shown in figures were generated by averaging the PC1 and PC2 coordinates across animals within each group at each successive time point. Display-only smoothing was applied to plotted group mean trajectories, while all quantitative analyses were performed on the unsmoothed trajectories.

The similarity of each AgRP axon stimulation mouse to the AgRP neuron cell body response was quantified using Euclidean distance in PC1-PC2 space. For each AgRP cell body mouse, leave-one-out reference trajectories were used so that each mouse was compared with the mean trajectory of the remaining mice. Then, at each time point, the distance between each AgRP axon stimulation mouse’s trajectory and the mean AgRP neuron cell body stimulation trajectory was calculated and then averaged across the full analysis window to yield a single distance value per animal, with smaller values indicating a more AgRP cell body-like response profile. Group-level descriptive similarity was also calculated from the mean distance between each group trajectory and the AgRP mean trajectory across time.

#### Exclusion criteria

After experimentation, brain regions of interest were manually inspected in every control and experimental mouse for viral injection and/or fiber placement. Mice were excluded from data analysis for insufficient or absent viral expression in the intended target region or off-target photometry fiber, optogenetic fiber, or cannula placement. Mice were also excluded if a photometry or optogenetic fiber became dislodged during the experiment.

#### Behavioral procedures

##### General behavioral procedures

Unless otherwise stated, mice were tested under ad libitum-fed conditions during the light phase of the light/dark cycle. Behavioral testing was performed at approximately the same time of day across repeated sessions. Experimental conditions were tested using counterbalanced designs, including drug versus vehicle and optogenetic light ON versus OFF conditions, with repeated test sessions separated by at least 24-48 h.

Before experimental testing, mice underwent three habituation sessions on separate days. Habituation included handling, i.p. injections, placement in the testing room and testing cage with food, and attachment to the fiber photometry and/or optogenetic patch cords used during subsequent testing. All implants were cleaned with 70% ethanol and connected to the photometry and/or optogenetic patch cords before mice were placed into a clean cage identical to their home cage and containing fresh bedding and access to water. Recordings began within several minutes of placement in the test cage.

For experiments involving chow presentation, a pre-weighed 3-5 g piece of standard laboratory chow (Purina 5001) was placed into the cage approximately 2-3 inches in front of the mouse. This standardized presentation procedure was used across all experiments involving chow.

Food intake was determined by weighing the remaining food and collecting and weighing crumbs generated during the session. Crumb weight was subtracted when calculating consumption.

##### Chow presentation during AgRP neuron manipulation

For chemogenetic activation or inhibition of AgRP neurons, recording began at session onset and mice received CNO (1 mg/kg, i.p.) or vehicle 15 min later. Chow was presented 45 min after session onset and food intake was measured at 75 min.

For optogenetic excitation of AgRP cell bodies or AgRP axon terminals, optical stimulation began 15 min after session onset using the stimulation parameters described above and continued for the remainder of the session. Chow was presented at 30 min and intake was measured at 75 min.

##### Object presentation

A yellow 1.5-mL Eppendorf tube was used as the test object. Mice were habituated to this object in their home cage beginning the night before testing and the objects remained in their home cages throughout the testing period. During experimental sessions, the object was placed 2-3 inches in front of the mouse. For chemogenetic experiments, CNO or vehicle was administered 15 min after session onset and the object was presented at 45 min. Recordings continued until 60 min. For optogenetic experiments, optical stimulation began at 15 min, the object was presented at 30 min, and the session ended at 45 min.

##### Novel food presentation

For novel food experiments, mice received CNO or vehicle 15 min after session onset. 30 min later, approximately 2 g of a novel chocolate-flavored grain pellet (TestDiet 5TUL, 181123) were presented in a small plastic weigh boat to which mice had previously been habituated. Intake was measured at 75 min using the same procedure described for chow, including collection and weighing of crumbs. Mice that failed to consume the novel food during the initial test were provided 2-4 h of access to the same pellets in their home cage on the following day. We confirmed that all mice consumed the pellets between the first and second test sessions. All mice were subsequently retested 48 h after the first test using an identical experimental procedure, such that the previously novel food was familiar at the time of the second test.

##### Cocaine administration

For experiments examining dopamine responses to cocaine during AgRP neuron activation, mice received CNO or vehicle 15 min after session onset in a counterbalanced design. Cocaine (5 mg/kg, i.p.) was administered to all mice 45 min after session onset, and dopamine signals were recorded until 75 min.

##### NAc dopamine receptor antagonism

For NAc pharmacology experiments, mice were connected to an intracranial infusion line connected to a syringe in a Harvard Apparatus syringe pump. Vehicle or flupentixol (12 or 20 µg) was infused unilaterally into the NAc in a volume of 350 nL at a rate of 0.3 µL/min. Infusions began at session onset. Immediately following the infusion, mice received i.p. CNO or vehicle. Chow was presented 30 min after the CNO injection, and food intake was measured at 60 min.

##### Optogenetic inhibition of VTA dopamine neurons

For experiments examining the contribution of VTA dopamine neurons to AgRP neuron-driven feeding, DAT-Cre;Npy-ires2-FlpO-D mice expressing Cre-dependent eNpHR3.0 in VTA dopamine neurons and Flp-dependent hM3Dq in NPY/AgRP neurons were tethered to the photometry patch cord and bilateral optogenetic patch cords. Mice received CNO or vehicle 15 min after session onset to activate AgRP neurons. Chow was presented at 45 min. Continuous 590-nm illumination was delivered from 44 min to 50 min or from 54 min to 60 min (so that VTA dopamine neurons were inhibited for about 5 min surrounding chow presentation, or 10 min later), depending on the experimental condition. Food intake was measured at 75 min. For closed-loop inhibition sessions, mice were either given a blue (450-nm) light stimulation (as a control) or orange (590-nm) light stimulation for inhibition. Mice received CNO or vehicle 15 min after session onset to activate AgRP neurons. Chow was presented at 45 min and final food intake was measured at 75 minutes. Each time a mouse approached within a 2-inch radius of the food pellet, the laser light was turned on in a 1s ON 1s OFF pattern that stayed on as long as the mouse was in the food zone.

##### Intra-PVH pharmacology

For PVH pharmacology experiments, mice were tethered to the photometry patch cord and an intracranial infusion line connected to a Harvard Apparatus syringe pump and secured to the implanted cannula with a fixing screw (RWD Life Science, model 62502). NPY (1 µg), SHU9119 (37 pmol), baclofen (20 ng), or vehicle (PBS) was infused unilaterally into the PVH in a volume of 200 nL at a rate of 0.2 µL/min. Infusions began following a 15-min baseline period and required approximately 1 min to complete. Chow was presented at 30 min and food intake was measured at 60 min.

##### Optogenetic inhibition of AgRP terminals in the PVH

To determine the contribution of AgRP projections to the PVH during AgRP neuron activation, AgRP-Cre x Ai39 mice [expressing both eNpHR3.0 (genetically) and hM3Dq (virally) in AgRP neurons] were tethered to a photometry patch cord recording from the NAc and an optogenetic patch cord targeting the PVH. At 15 min after session onset, mice received CNO in both conditions, and continuous 590-nm illumination of AgRP terminals in the PVH was initiated during light ON sessions. The order of light ON and light OFF conditions was counterbalanced across mice. Illumination continued for the remainder of the session. Chow was presented at 30 min and food intake was measured at 75 min.

##### Chemogenetic inhibition of PVH Y1R neurons

For photometry experiments examining the effect of PVH Y1R neurons on dopamine signaling, mice expressing hM4Di in PVH Y1R neurons were tethered to the photometry patch cord and received CNO or vehicle 15 min after session onset. Chow was presented at 45 min and food intake was measured at 75 min. In a separate feeding experiment, mice received CNO or vehicle and were provided ad libitum access to chow. Food intake was measured hourly for 6 h.

#### Slice electrophysiology

Npy-ires2-FlpO-D x Dat-Cre mice underwent stereotaxic surgery at approximately P28. Experimental mice received a 300-nL injection of AAV8-hSyn-fDIO(Gq)-mCherry-WPREpA (Addgene, #154868-AAV8) into the ARC (-0.90 mm AP, 0.20 mm ML, -5.90 and -5.75 mm DV) to express hM3Dq selectively in AgRP/NPY neurons. AAV9-Syn-DIO-eGFP (Addgene, #50457-AAV9) was injected into the VTA (-2.80 mm AP, 0.50 mm ML, -4.40 mm DV; 400nL) to identify DAT-expressing dopamine neurons in subsequent electrophysiological recordings. Control mice were DAT-Cre mice that received the same Cre-dependent eGFP injection into the VTA. Before slice electrophysiology experiments, chemogenetic functionality (stimulation of AgRP/NPY neurons) was behaviorally validated. Experimental and control mice received i.p. CNO (1 mg/kg) and food intake was measured after one hour. We verified increased food intake in experimental relative to control mice to confirm functional activation of NPY/AgRP neurons (**Figure 1H**).

##### Acute brain slice preparation

At P45-50, mice were euthanized and acute brain slices containing the VTA were prepared for whole-cell electrophysiological recordings from eGFP-positive DAT-expressing neurons. All mice were sacrificed between 10 am and 11 am (ZT 3-4). The brain was rapidly removed and transferred to an ice-cold (∼4°C), oxygenated solution containing, in mM: 124 choline chloride, 2.5 KCl, 1.23 NaH2PO4, 26 NaHCO3, 1 CaCl2, 6 MgCl2, and 10 glucose (pH 7.3 adjusted with NaOH, 300–310 mOsm).

Acute horizontal midbrain slices containing the VTA were prepared at a thickness of 300 µm using a vibratome (Vibratome 1000). Slices were immediately transferred to a holding chamber containing artificial cerebrospinal fluid (aCSF) continuously bubbled with 95% O2, 5% CO2 and maintained at room temperature (22-24 °C) for at least 1 h before recording. The standard aCSF contained (in mM): 124 NaCl, 2.5 KCl, 1.2 NaH2PO4,24 NaHCO3, 10 glucose, 2 CaCl2, and 1.3 MgCl2 (pH 7.3–7.4, 295-305 mOsm).

##### Visual identification and electrophysiological confirmation of VTA dopamine neurons

Slices were transferred to a submerged recording chamber and continuously perfused with oxygenated aCSF at a flow rate of 1-2 mL/min at 32-34 °C. Dopaminergic neurons within the VTA were visually targeted by their eGFP fluorescence using an upright microscope equipped with IR-DIC optics and a FITC filter set.

Identified eGFP neurons were further electrophysiologically validated by assessing key dopaminergic properties. In voltage-clamp mode, hyperpolarizing voltage steps (from -50 mV to -110 mV in -10 mV increments) were applied to evoke a characteristic hyperpolarization-activated cation current (*I*h). In cell-attached or current-clamp mode, target cells exhibited a broad action potential waveform and a low baseline firing rate (<5 Hz).

##### Whole-cell patch-clamp recordings of mEPSCs and mIPSCs

Whole-cell patch-clamp recordings were obtained using borosilicate glass pipettes (2–4 MΩ resistance) pulled with a horizontal micropipette puller. Recording electrodes were filled with Cs+ based internal solution containing (in mM): 135 cesium methanesulfonate, 2 MgCl2, 10 HEPES, 1.1 EGTA, 2 Mg-ATP, 10 Na2-phosphocreatine, and 0.3 Na2-GTP (pH 7.3 adjusted with CsOH, 285–290 mOsm).

Miniature postsynaptic currents were recorded in the continuous presence of tetrodotoxin (TTX; 0.5 µM) to block action-potential-dependent synaptic transmission. Miniature excitatory postsynaptic currents (mEPSCs) were recorded as inward currents at a holding potential of -60 mV, whereas miniature inhibitory postsynaptic currents (mIPSCs) were recorded as outward currents at a holding potential of 0 mV.

The pharmacological identities of the recorded events were confirmed in recordings with receptor antagonists. Inward miniature currents recorded at -60 mV were blocked by combined application of the AMPA/kainate receptor antagonist CNQX (10 µM) and the NMDA receptor antagonist APV (50 µM), confirming their identity as glutamatergic mEPSCs. Outward miniature currents recorded at 0 mV were blocked by the GABAA receptor antagonist bicuculline (30 µM), confirming their identity as mIPSCs

##### Data acquisition and analysis

Data were acquired using a MultiClamp 700B amplifier (Molecular Devices) controlled by AxoGraph X software (AxoGraph Scientific). Signals were low pass filtered at 2.4 kHz and digitized at 10 kHz. Series resistance Rs was continuously monitored throughout the experiment. Cells were retained when series resistance was ≤20 MOhm both before and after recording and changed by ≤20%.

For mIPSC and mEPSCs analyses, events were detected offline in AxoGraph using template matching. For mIPSCs, recordings were held at 0 mV and detected using a template with a 5-ms baseline, 10-ms template duration, +30-pA amplitude, 0.2-ms rise time, 10-ms decay time, 15-ms analysis length, and signal-to-noise threshold of 4. For mEPSCs, recordings were held at -60 mV, the detection template used a minimum inter-event separation of 3 ms, template amplitude of -20 pA, rise time of 0.2 ms, decay time of 5 ms, and a signal-to-noise threshold of -3.5; events within a diagnostic amplitude range of 15-400 pA were retained. Detected event counts and the actual recording duration were used to calculate cell level frequency, and the median event amplitude within each cell was used for amplitude analyses. Statistical significance was calculated using a two-sided Mann-Whitney U test between groups. In addition, because cells from the same mouse are not independent, a linear mixed-effects model was fitted to log10-transformed cell values, with group as a fixed effect and mouse as a random intercept:

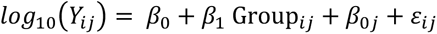

where *Y_i_*_j_ is frequency or median amplitude for cell *i* from mouse *j*; Group is coded 0 for control and 1 for test; *β_0j_* is the mouse-specific random intercept; and *ε_ij_* is the residual error.

Log10 transformation was used to stabilize variance. Model estimates were back-transformed and reported as test/control ratios with 95% confidence intervals and two-sided p-values.

#### Histology and imaging

##### Verification of viral expression and fiber placement

Mice were deeply anesthetized with isoflurane and transcardially perfused with 0.1 M phosphate-buffered saline (PBS), followed by 4% paraformaldehyde (PFA). Brains were removed and post-fixed in 4% PFA at 4°C for at least 4 h before transfer to PBS. For routine verification of viral expression and fiber or cannula placement, 100-µm coronal sections were prepared using a vibrating blade microtome (Leica VT1000S). Epifluorescence images were acquired using a Keyence BZ-X800 slide-scanning microscope to verify viral expression and implant placement.

##### RNA in situ hybridization

For RNAscope experiments, brains were collected from AgRP-ires-Cre mice that received hM3Dq viral injections and from uninjected controls. Following perfusion, brains were postfixed overnight in 4% PFA, cryoprotected in 30% sucrose for 24-72 h, frozen at -80 °C, and sectioned on a cryostat (Leica) at 16 µm. Sections were mounted directly onto slides and stored at -80 °C until processing. RNA in situ hybridization was performed using the RNAscope Multiplex Fluorescent Detection Kit v2 according to the manufacturer’s instructions with probes and reagents obtained from Advanced Cell Diagnostics. Sections were washed to remove OCT, postfixed in 4% PFA for 15 min, dehydrated through graded ethanol solutions, treated with hydrogen peroxide, subjected to target retrieval at 95 °C for 5 min, and incubated with Protease III at 40°C for 30 min. Sections were hybridized with probes targeting *Fos* and *Npy1r* for 2 h at 40°C, followed by sequential AMP1, AMP2, and AMP3 amplification steps. Signals were developed sequentially using HRP-mediated amplification with Cy5 and Cy3 Opal fluorophores diluted 1:1500 in TSA buffer, with HRP blocking performed between channels. Following the final wash, sections were coverslipped with VectaShield+DAPI (Vector Laboratories, H-1200-10) and Mowiol (Sigma-Aldrich, # 81381).

Epifluorescence images were acquired using a BZ-X800 slide scanner (Keyence) with 2X, 4X, 10X, and 20X objective lenses, and confocal images were acquired using a Stellaris 5 microscope (Leica, purchased with S10OD030354) with a 20X objective lens. Two to three sections per mouse were analyzed for each experiment. Images were analyzed in Fiji/ImageJ, and DAPI, *Fos*, and *Npy1r* cells were quantified.

#### Statistical analyses

Data are presented as mean ± SEM unless otherwise indicated. Statistical analyses were performed using Prism 11 (GraphPad Software). Paired two-tailed t-tests were used for within-subject comparisons and unpaired two-tailed t-tests were used for between-group comparisons. Repeated-measures ANOVAs were used for experiments containing three or more within-subject conditions, one-way ANOVAs were used for comparisons among independent groups, and two-way ANOVAs were used to assess the effects of treatment and time. Repeated-measures ANOVAs were performed with Geisser-Greenhouse correction for sphericity. Post hoc multiple comparisons were performed using Tukey’s, Šídák’s, or Dunnett’s tests as appropriate. Pearson correlations were used to assess relationships between continuous variables. For PCA analyses, mouse-level Euclidean distances from the AgRP trajectory were compared across groups using one-way ANOVA followed by Dunnett’s multiple comparisons test with the AgRP cell-body stimulation group as the reference. Statistical outliers were identified using Grubbs’ test (*p* < 0.05) and excluded from subsequent analyses: across all experiments, one mouse each was excluded from Figure 1E, 1F, 4E,H,I, and 5C. Statistical significance was defined as *p* < 0.05. Sample sizes, statistical tests, test statistics, and multiple-comparison procedures for individual experiments are provided in Table S1.

## Notes

### Summary of Updates

Data and statistics have been updated

